# An Ultra-High-Resolution 17.2 T MRI Atlas (HypoAtlas) and Multimodal Pipeline to Study the Mouse Hypothalamus: Sexual Dimorphism and Lateralization Insights

**DOI:** 10.64898/2026.01.08.698542

**Authors:** Alicia Sicardi, Pierre Labouré-Santavicca, Ivy Uszynski, Fawzi Boumezbeur, Cyril Poupon, Philippe Vernier, Philippe Ciofi, Pierre-Yves Risold, Vincent Prévot, Luisa Ciobanu

## Abstract

Structural and functional insights into the mouse hypothalamus are hampered by its small size and deep location. Here, we leverage ultra-high-field magnetic resonance imaging (UHF-MRI) at 17.2 Tesla to achieve unprecedented spatial resolution in structural, functional and neurochemical imaging of the mouse hypothalamus, including sexual dimorphism in certain nuclei. High-resolution *ex vivo* anatomical MRI enabled precise hypothalamic parcellation, improving on existing atlases and revealing nuclei previously unresolved by MRI. Diffusion MRI and tractography mapped intra- and extra-hypothalamic pathways, facilitating circuit-level exploration without *a priori* assumptions. Resting-state fMRI combined with independent component analysis identified novel hypothalamic networks, demonstrating the enhanced capacity of UHF MRI to detect deep-brain activity. ¹H-magnetic resonance spectroscopy quantified neurochemical profiles, revealing sexually dimorphic heterogeneity within the hypothalamus. Our comprehensive multimodal approach uncovers sex differences in hypothalamic anatomy, microstructure, and neurochemistry, emphasizing the importance of sex as a biological variable. This integrated pipeline offers a valuable resource for dissecting hypothalamic circuits and functions, advancing our understanding of neuroendocrine regulation, behavior, and disease mechanisms, with direct translational relevance.

## Introduction

The hypothalamus, a small structure located at the base of the brain, exhibits one of the highest levels of neuronal diversity and projection pattern complexity among brain regions ^1, 2^. It functions as the central regulator of numerous vital processes, including growth, reproduction, energy homeostasis, feeding behavior, thirst and fluid balance, circadian rhythms, sleep and social interactions. The highly specialized neuronal and glial populations of these nuclei communicate both physically and functionally to integrate and relay information related to external and internal environmental cues to extrahypothalamic structures, allowing the brain to maintain body homeostasis and coordinate appropriate adaptive behavioral responses essential for species survival^3^. As such, the study of the hypothalamus is fundamental to understanding many of the most prevalent and life-threatening disorders in modern society, including obesity, metabolic syndrome, eating disorders, sleep disturbances, reproductive dysfunction, cognitive impairments, psychiatric disorders and neurodegenerative diseases^3–6^.

Fundamental and preclinical research is largely focused on the mouse model, *Mus musculus*, due to the availability of a fully characterized genome^7^, the ability to visualize and manipulate gene expression, including in the brain^8,9^, and its ease of reproduction and small size, ideal for routine and cost-effective husbandry. However, despite significant progress in phenotyping hypothalamic cell types in mice thanks to advances in single-cell RNA sequencing and spatial transcriptomics^10–12^, most of our neuroanatomical knowledge of the hypothalamus has been derived from studies in the rat, *Rattus norvegicus*^2, 13–15^. Both from the point of view of advancing our knowledge of how this key structure works, and to develop targeted pharmacological and genetic therapies for disorders involving its dysfunction, it is thus crucial to acquire a comprehensive understanding of hypothalamic structure, function and connectivity in the mouse, including how these differ throughout the lifespan and across sexes.

Classic neuroanatomical studies predominantly utilize techniques such as large-scale *in situ* hybridization, viral-, tract-tracing and immunohistochemistry combined with light microscopy^16, 17^. These methods enable the visualization of neuronal fiber organization, the identification of neuronal populations within structures, and transcriptomic profiling, providing detailed insights into cellular and circuit architecture. In addition, fluorescence micro-optical sectioning tomography has been used to reconstruct projections from single labeled axons from multiple peptidergic neuron populations in the hypothalamus^18^. While these approaches are highly detailed and specific, they are often time-consuming, typically analyzed after the animals are sacrificed, and require the integration of multiple tools to assemble comprehensive datasets. Additionally, they may not fully represent *in vivo* or overall brain organization, nor allow for longitudinal studies, which limits their direct translatability to human research.

Over the past two decades, magnetic resonance imaging (MRI) has emerged as an indispensable *in vivo* brain imaging technique to map brain structure and connectivity in rodent models. Originally developed for the non-invasive imaging of the human brain, MRI has benefited from significant technological advances, such as increased magnetic field strength and improved coil designs, that now permit the imaging of rodents at the submillimeter scale^19^. The non-invasive and painless nature of MRI not only adheres to ethical principles in animal research but also enhances its translational potential for human patients. However, the spatial resolution and tissue specificity of traditional MRI do not yet match those of conventional histology. Thus, in contrast to human studies, which have successfully generated detailed MRI atlases of specific hypothalamic nuclei and begun to explore their structural connectivity^20, 21^, the limited resolution of rodent MRI with respect to the size of the hypothalamus generally results in its being segmented only as a whole or into broad subregions, without the precise delineation of individual nuclei^19^.

Recent breakthroughs in ultra-high-field MRI—particularly for rodent models—have dramatically advanced brain imaging, offering unparalleled resolution and detail^22–24^. In this study, we used a 17.2 Tesla (T) ultra-high field (UHF) MRI system to achieve highly detailed imaging of the mouse hypothalamus and its distinct nuclei, with a resolution approaching that of histology. This capability allowed for a comprehensive and multimodal exploration of hypothalamic nuclei, focusing on their structural and functional connectivity, while also highlighting sexual dimorphism in the organization and microstructure of individual nuclei. Additionally, 17.2 T MRI generated spectra of unprecedented quality through proton-magnetic resonance (¹H-MR) spectroscopy that enabled us to pinpoint real-time, sex-dependent metabolic activity and dynamic interactions within the hypothalamus, potentially facilitating *in vivo* studies of human brain networks and opening new horizons for neuroscience research.

## Results

### Segmentation of Mouse Hypothalamic Nuclei in 40-µm resolution MRI space

A 40-µm isotropic resolution mouse brain template was generated based on twelve three-dimensional (3D) T_2_-weighted images acquired *ex vivo* at 17.2 T. This template was then used to manually segment hypothalamic nuclei, identified visually based on anatomical landmarks derived from histological cross-sections, with additional guidance from previously published atlases, including the DSURQE MRI atlas ^25–27^ and the Allen Mouse Brain Atlas. MRI atlas planes were aligned with the coronal plan levels described in the Paxinos and Franklin atlas^28^ (Figure 1a-1f). By systematically segmenting visible hypothalamic nuclei across MRI sections in the coronal, sagittal (Figure 1g) and horizontal (Figure 1h) planes within the whole-brain MRI space, we delineated the 3D structure of each nucleus. This multi-planar approach enabled the construction of a detailed 3D representation of hypothalamic nuclei (Figure 1i), as follows.

**Figure 1.**
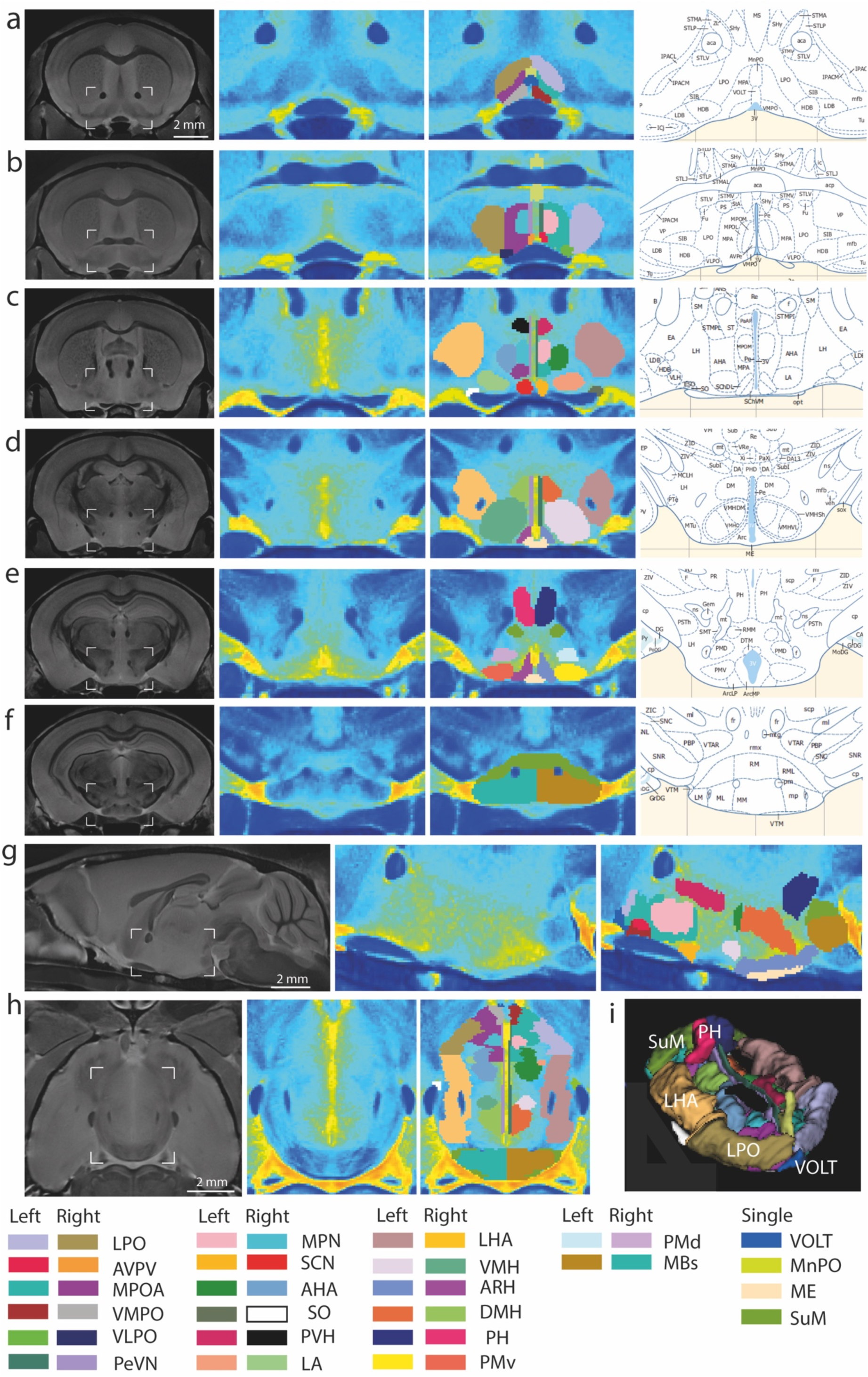
Segmentation of Mouse Hypothalamic Nuclei. Visualization of the mouse brain in grayscale, highlighting hypothalamic regions in coronal plane (Column 1, **a-f**). Zoomed views of the hypothalamic region without segmentation of the nuclei are shown in Column 2. Segmented hypothalamic nuclei are displayed in Column 3, with distinct colors used to differentiate right and left hemispheres. Column 4 shows the corresponding regions in the Paxinos and Franklin reference atlas. The regions illustrated include (**a, b**) the preoptic area, (**c**) the anterior hypothalamic area, (**d**) the tuberal hypothalamic area, (**e**) the transition between the tuberal and posterior hypothalamic areas, and (**f**) the posterior hypothalamic area Additional views include (**g**) a sagittal section, (**h**) a horizontal section, and (**i**) a three-dimensional rendering of all segmented nuclei. Abbreviations are listed in Table 1.

**Table 1.**
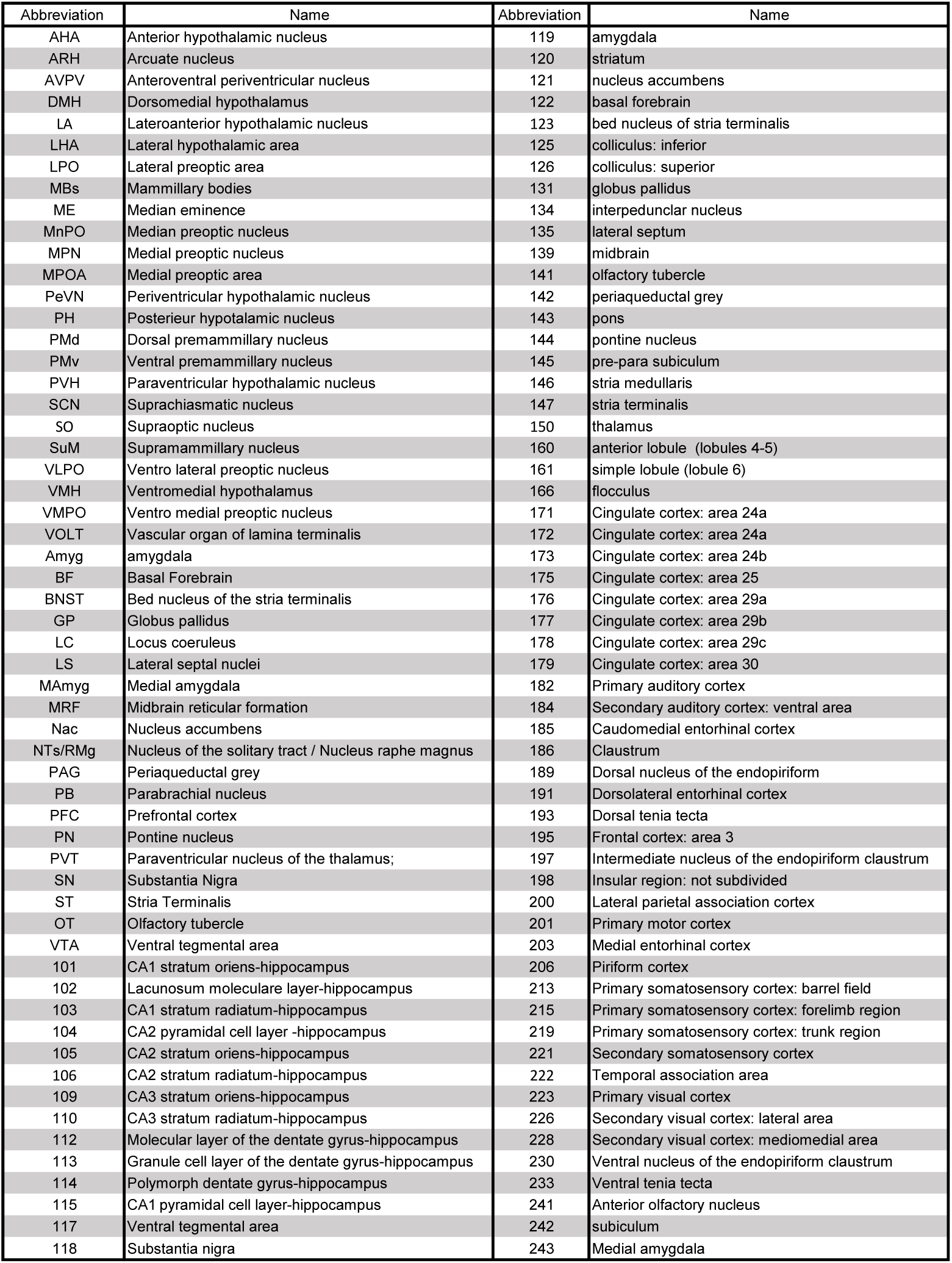
List of abbreviations and numerical correspondences for the hypothalamic nuclei and areas used and present in the analyses.

| Abbreviation | Name | Abbreviation | Name |
| --- | --- | --- | --- |
| AHA | Anterior hypothalamic nucleus | 119 | amygdala |
| ARH | Arcuate nucleus | 120 | striatum |
| AVPV | Anteroventral periventricular nucleus | 121 | nucleus accumbens |
| DMH | Dorsomedial hypothalamus | 122 | basal forebrain |
| LA | Lateroanterior hypothalamic nucleus | 123 | bed nucleus of stria terminalis |
| LHA | Lateral hypothalamic area | 125 | colliculus: inferior |
| LPO | Lateral preoptic area | 126 | colliculus: superior |
| MBs | Mammillary bodies | 131 | globus pallidus |
| ME | Median eminence | 134 | interpeduncular nucleus |
| MnPO | Median preoptic nucleus | 135 | lateral septum |
| MPN | Medial preoptic nucleus | 139 | midbrain |
| MPOA | Medial preoptic area | 141 | olfactory tubercle |
| PeVN | Periventricular hypothalamic nucleus | 142 | periaqueductal grey |
| PH | Posterior hypothalamic nucleus | 143 | pons |
| PMd | Dorsal premammillary nucleus | 144 | pontine nucleus |
| PMv | Ventral premammillary nucleus | 145 | pre-para subiculum |
| PVH | Paraventricular hypothalamic nucleus | 146 | stria medullaris |
| SCN | Suprachiasmatic nucleus | 147 | stria terminalis |
| SO | Supraoptic nucleus | 150 | thalamus |
| SuM | Supramammillary nucleus | 160 | anterior lobule (lobules 4-5) |
| VLPO | Vento lateral preoptic nucleus | 161 | simple lobule (lobule 6) |
| VMH | Ventromedial hypothalamus | 166 | flocculus |
| VMPO | Vento medial preoptic nucleus | 171 | Cingulate cortex: area 24a |
| VOLT | Vascular organ of lamina terminalis | 172 | Cingulate cortex: area 24a |
| Amyg | amygdala | 173 | Cingulate cortex: area 24b |
| BF | Basal Forebrain | 175 | Cingulate cortex: area 25 |
| BNST | Bed nucleus of the stria terminalis | 176 | Cingulate cortex: area 29a |
| GP | Globus pallidus | 177 | Cingulate cortex: area 29b |
| LC | Locus coeruleus | 178 | Cingulate cortex: area 29c |
| LS | Lateral septal nuclei | 179 | Cingulate cortex: area 30 |
| MAmyg | Medial amygdala | 182 | Primary auditory cortex |
| MRF | Midbrain reticular formation | 184 | Secondary auditory cortex: ventral area |
| Nac | Nucleus accumbens | 185 | Caudomedial entorhinal cortex |
| NTs/RMg | Nucleus of the solitary tract / Nucleus raphe magnus | 186 | Clastrum |
| PAG | Periaqueductal grey | 189 | Dorsal nucleus of the endopiriform |
| PB | Parabrachial nucleus | 191 | Dorsolateral entorhinal cortex |
| PFC | Prefrontal cortex | 193 | Dorsal tenia tecta |
| PN | Pontine nucleus | 195 | Frontal cortex: area 3 |
| PVT | Paraventricular nucleus of the thalamus; | 197 | Intermediate nucleus of the endopiriform claustrum |
| SN | Substantia Nigra | 198 | Insular region: not subdivided |
| ST | Stria Terminalis | 200 | Lateral parietal association cortex |
| OT | Olfactory tubercle | 201 | Primary motor cortex |
| VTA | Ventral tegmental area | 203 | Medial entorhinal cortex |
| 101 | CA1 stratum oriens-hippocampus | 206 | Piriform cortex |
| 102 | Lacunosum moleculare layer-hippocampus | 213 | Primary somatosensory cortex: barrel field |
| 103 | CA1 stratum radiatum-hippocampus | 215 | Primary somatosensory cortex: forelimb region |
| 104 | CA2 pyramidal cell layer -hippocampus | 219 | Primary somatosensory cortex: trunk region |
| 105 | CA2 stratum oriens-hippocampus | 221 | Secondary somatosensory cortex |
| 106 | CA2 stratum radiatum-hippocampus | 222 | Temporal association area |
| 109 | CA3 stratum oriens-hippocampus | 223 | Primary visual cortex |
| 110 | CA3 stratum radiatum-hippocampus | 226 | Secondary visual cortex: lateral area |
| 112 | Molecular layer of the dentate gyrus-hippocampus | 228 | Secondary visual cortex: mediomedial area |
| 113 | Granule cell layer of the dentate gyrus-hippocampus | 230 | Ventral nucleus of the endopiriform claustrum |
| 114 | Polymorph dentate gyrus-hippocampus | 233 | Ventral tenia tecta |
| 115 | CA1 pyramidal cell layer-hippocampus | 241 | Anterior olfactory nucleus |
| 117 | Ventral tegmental area | 242 | subiculum |
| 118 | Substantia nigra | 243 | Medial amygdala |

#### Preoptic region

The most anterior region considered here is the preoptic region (Figure 1a-1b). This region constitutes the periventricular gray of the most rostral part of the third ventricle (3V). It comprises multiple nuclei and controls many social behaviors and homeostatic functions^9^, including sleep^29^. It is located on either side of the anterior 3V, between the anterior commissures and the optic chiasma (Figure 1a, 1b). At its anterior extremity, the 3V is capped dorsally by the organum vasculosum laminae terminalis (VOLT) ^30^, which is itself covered by the median preoptic nucleus (MnPO) on the dorsal midline (Figure 1a). The OVLT and MnPO play key roles in the regulation of water intake^31, 32^ and body temperature^33^, and contain most of the hypothalamic GnRH-expressing neurons that control reproduction^34^, as well as *Nos1*-expressing neurons that modulate their activity^35–37^. On each side of these structures lies the medial preoptic area (MPOA), involved in parenting and mating, a process that appears to involve, at least in part, galaninergic neurons^38^ (Figure 1a). The MPOA is bordered dorsally (Figure 1a) and laterally (Figure 1b) by the lateral preoptic area (LPO) and ventrolaterally by the ventrolateral preoptic nucleus (VLPO) (Figure 1a, 1b). While the LPO appears to be involved in nociceptive processes^39^, the VLPO is known to play a role in sleep regulation^40, 41^. Ventromedially to the MPOA and on each side of the emerging 3V sits the ventromedial preoptic area (VMPO) containing *Pacap*-expressing neurons involved in the regulation of body temperature^42^. Embedded in the more caudal aspect of the MPOA is the medial preoptic nucleus (MPN) (Figure 1b), which is involved in the control of social behavior^43^ and contains subgroups of neuronal populations that are sexually dimorphic in rats, with more neurons in males than in females^44^. Among sexually dimorphic nuclei, the anteroventral periventricular nucleus (AVPV) is located ventrally between the wall of the3V and the MPOA (Figure 1b). The AVPV plays a crucial role in promoting the preovulatory gonadotropin surge through its Kiss1 neuronal population^44, 45^.

#### Anterior region

Caudally, beyond the decussation of the anterior commissure—which delineates the posterior boundary of the preoptic region and the penetration of the fornix into the diencephalon — the MPOA transitions into the anterior hypothalamic area (AHA) (Figure 1c), which contains cell populations involved in the control of cardiovascular function^46^. Between the AHA and the 3V lies the periventricular nucleus (PeVN) (Figure 1c), which contains A14 dopaminergic neurons involved in the regulation of either locomotion or prolactin secretion, depending on their projection targets^47^. Ventral to the PeVN on each side of the ventral edge of the 3V lies the suprachiasmatic nucleus (SCN) (Figure 1c), the seat of the master circadian clock^48^. Bordering the dorsal edge of the 3V on each side is the paraventricular nucleus of the hypothalamus (PVH) (Figure 1c), a key hypothalamic hub that coordinates autonomic, neuroendocrine and behavioral functions^49^. The mouse PVH has its first cells at the entrance of the fornix into the diencephalon, in a position slightly ventral to this tract. It begins to emerge as a distinct, slim, cylindrical nucleus at the midrostral to caudal level of the anterior hypothalamic nucleus, where the dense distribution of magnocellular and parvocellular neurons makes the nucleus highly visible. Caudal to this level, the entire PVH, particularly its dorsal portion, extends laterally and appears as a triangular nucleus with its medial edge parallel to, and its dorsal edge perpendicular to the wall of the 3V. At its most caudal level, the PVH forms a lateral wing that extends through the lateral hypothalamic area (LHA), terminating above the descending column of the fornix. The LHA, which contains neuronal populations involved in the regulation of motivated behaviors, feeding, energy balance and sleep^50, 51^, lies at the lateral outer border of the AHA (Figure 1c), marking a transition from the LPO (Figure 1b). It should be noted that the LHA remains an extremely heterogeneous region of the brain in terms of its neuronal populations and is still poorly understood ^52^. Its anatomical boundaries are not well established and are still being studied ^52^. Lying along the most lateral border of the optic tracts bilaterally is the supraoptic nucleus (SO), predominantly composed of neuroendocrine vasopressin- and oxytocin-expressing neurons —also present in the PVH—that project to the posterior pituitary to regulate osmoregulation and reproduction after the secretion of these hormones into the general circulation (Figure 1c) ^53–55^.

#### Tuberal region

The tuberal region of the hypothalamus begins with the emergence of the ventromedial nucleus of the hypothalamus (VMH), which is one of the largest hypothalamic structures readily identifiable in the T_2_-weighted 3D image in coronal sections (Figure 1d). The VMH appears as a clearly defined, inclined oval shape, situated on both sides of the 3V, bordered by the arcuate nucleus of the hypothalamus (ARH) ventrally, and the dorsomedial nucleus of the hypothalamus (DMH) dorsally. The VMH, which has been widely defined as consisting of dorsomedial, central/core and ventrolateral cytoarchitectonic subdivisions, is primarily involved in the regulation of energy homeostasis as well as social behaviors such as sexual and aggressive interactions^56^. The DMH, located bilaterally along the upper third of the 3V and separated from its wall by the caudal extent of the PeVN (Figure 1d) is involved in a broad spectrum of physiological processes, including thermogenesis, regulation of blood pressure, food intake and circadian rhythms in mice^33, 57–61^. The ARH, flanking the most ventral part of the 3V (Figure 1d), appears coronally immediately caudal to the rostral extent of the VMH. It exhibits a characteristic horse-shoe shape in coronal sections, situated on each side of the 3V. As the median eminence (ME) develops (Figure 1d), the ARH divides into two bilateral crescent-shaped nuclei positioned beneath the VMH and, subsequently, beneath the DMH. The ARH runs along each side of the 3V until it gradually disappears at the transition between the tuberal and mammillary regions. The ARH serves as a central hub in integrating peripheral metabolic signals^5, 62^ extravasating from the fenestrated capillary vessels of the ME and shuttled, at least in part, by specialized ependymal glial cells called tanycytes^63, 64^, and plays a vital neuroendocrine role in regulating growth hormone and prolactin secretion by the pituitary. Additionally, the neuronal populations of the ARH interact with and tightly regulate neuroendocrine secretion by other neurons projecting to the ME, which release their neurohormones into the pituitary portal blood vessels to modulate the activity of endocrine cells in the anterior pituitary^65^.

#### Mammillary region

The mammillary region of the hypothalamus begins with the appearance of the ventral premammillary nucleus (PMv) (Figure 1e). Its rostral tip aligns with the most anterior aspect of the 3V recess and extends toward the caudal end of the VMH. The PMv is medially bordered by the ARH and the dorsal premammillary nucleus (PMd), ventrally by the ventral boundary of the hypothalamus, and dorsally by the fornix, the PMd, and the posterior hypothalamic nucleus (PH) (Figure 1e). Its shape is reminiscent of a crescent moon with the concavity facing dorsolaterally towards the fornix. Caudally, it adopts an oval form with a broader transverse axis, with its boundaries located rostral to the mammillary bodies (MBs) (Figure 1f). The PMv is a key node that integrates internal and external environmental cues to regulate neuroendocrine responses and social behaviors^66^. The PMd is a differentiation of the medial mammillary nucleus with afferents from the ventral tegmental nucleus and projections to the anterior thalamus. It has the same developmental origin as the thalamus. It is specialized in integrating threatening predator information from different sensory modalities^67^. The PH, which is involved in modulating hippocampal oscillatory activity, movement control, memory processing and arousal^68^, lies dorsal to the supramammillary nucleus (SuM) and the MBs (Figure 1e,1f) and limited laterally by the mammillothalamic tract. The SuM is another crucial hypothalamic hub for several behavioral and cognitive processes, including reward-seeking, exploration and social memory^69^. The MBs, historically associated with temporal and contextual memory through their role in the hippocampus–cortex–hippocampus circuit or Papez circuit — known to be mediated by the hypothalamus — are a complex of nuclei located on the ventral floor of the posterior hypothalamus, appearing as a pair of spherical protrusions on the underside of the brain^70^. All abbreviations are detailed in the Table T1.

#### Atlas integration and automated analysis

Segmentations of the hypothalamic nuclei (HypoAtlas) were integrated into the DSURQE MRI atlas^25–27^, resulting in a hypothalamus-enriched atlas that was then used for all subsequent analyses.

#### Sexual Dimorphism in the Volumes of Hypothalamic Nuclei

Automated segmentation of the hypothalamus was performed on a set of high-resolution 3D T_2_-weighted *ex vivo* 17.2 T MRI acquisitions from perfusion-fixed female (n = 7) and male (n = 7) wild-type C57BL/6J mice (Figure 2a). Consistent with existing literature on humans^71–73^, monkeys^72^, sheep^74^, rats^75^ and Japanese quails^76^, the MPN was observed to be larger in males than in females (0.055 ± 0.002 for males and 0.044 ± 0.002 for females, respectively; Mann-Whitney test : U = 0, p = 0.0012) (Figure 2b). Other hypothalamic nuclei also exhibited sexual dimorphism. In particular, the volumes of the PH, PeVN and PVH, normalized to the whole brain, were greater in males compared to females (0.055 ± 0.001 for males and 0.052 ± 0.001 for females, Mann-Whitney test: U = 5, p = 0.0221 for the PH; 0.061 ± 0.002 for males and 0.052 ± 0.002 for females, Mann-Whitney test: U = 6, p = 0.035 for the PeVN; 0.034 ± 0.002 for males and 0.026 ± 0.001 for females, Mann-Whitney test: U = 4, p = 0.0140 for the PVH) (Figure 2b).

**Figure 2.**
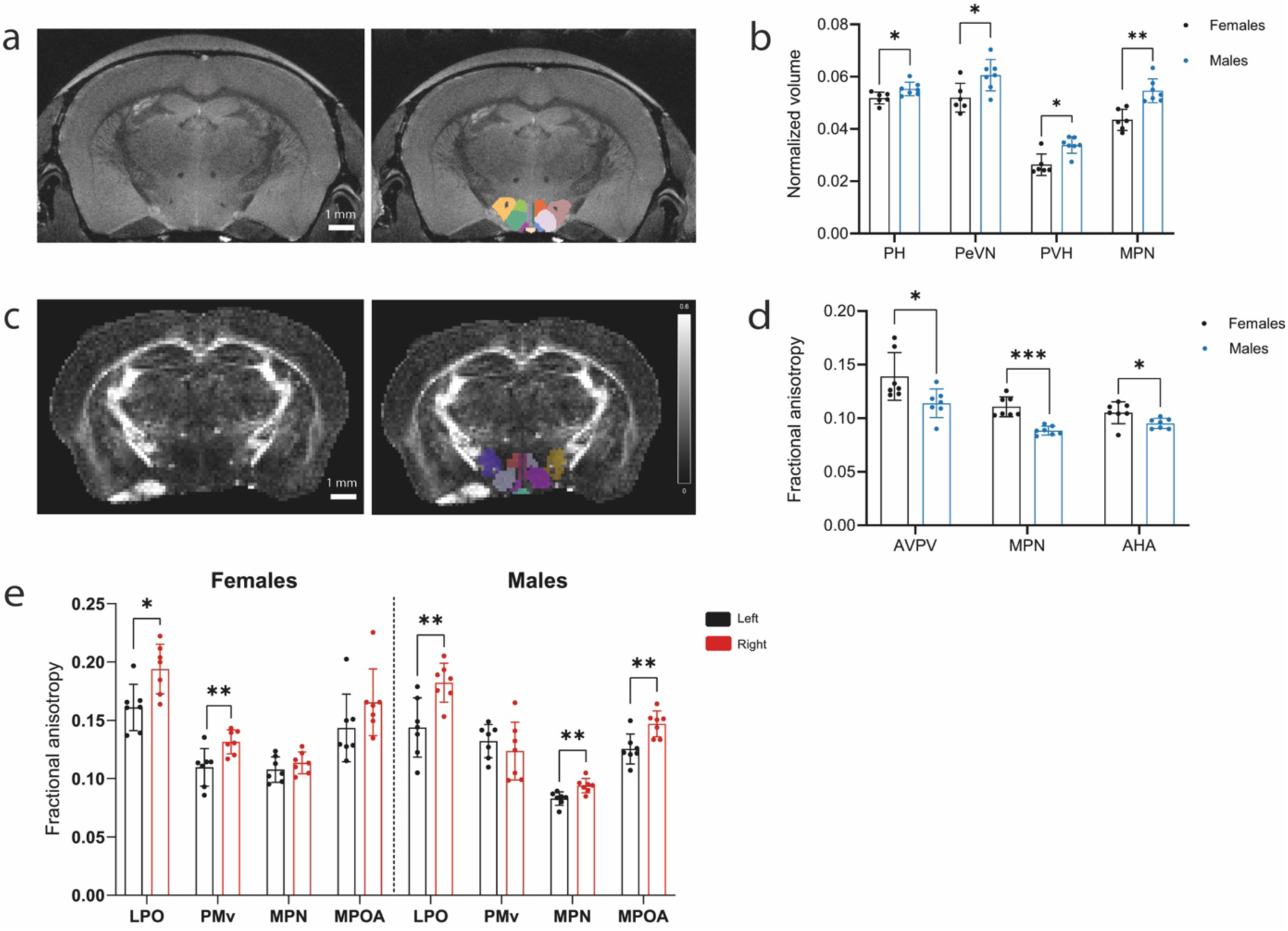
Sexual Dimorphism and Lateralization in Hypothalamic Nucleus Volumes and Diffusion-derived Metrics. (**a**) Illustration of an anatomical image acquired at 17.2 T without (left) and with (right) segmentation of the hypothalamic nuclei in coronal section at the level of the tuberal hypothalamus. (**b**) Bar graphs showing the volumes of sexually dimorphic hypothalamic nuclei normalized to the volume of the whole brain, with females shown in black and males in blue. Mann-Whitney U test, *p = 0.022, U = 5 for PH; *p = 0.035, U = 6 for PeVN; *p = 0.014, U = 4 for PVH; **p = 0.0012, U = 0 for MPN; n = 6, 7. (**c**) Fractional anisotropy (FA) map of a coronal section through the tuberal hypothalamus, with hypothalamic nuclei indicated; (**d**) Bar graphs representing the differences in FA between females and males. Mann-Whitney U test *p = 0.011, U = 5 for AVPV; ***p = 0.0006, U = 0, for MPN; *p = 0.026, U = 7 for AHA; n = 7 for both sexes. (**f**) Bar graphs representing the difference in FA between the right (red) and left (black) sides of the hypothalamus. For females, Mann-Whitney U test, *p = 0.011, U = 5 for LPO and **p = 0.007, U = 4 for PMv, n=7. For males, Mann-Whitney U test, **p = 0.007, U = 4 for LPO, **p = 0.0023, U = 2 for MPN and **p = 0.0041, U = 3 for MPOA, n = 7. Abbreviations: see Table 1.

No other hypothalamic nuclei studied showed sexual dimorphism in total volume. No lateralization was observed in any nucleus.

#### Sex Differences in MRI Diffusion Metrics of Hypothalamic Nuclei

Water molecule diffusion measurements provide insights into the tissue’s microenvironment, independent of volume. In this study, fractional anisotropy (FA), mean diffusivity (MD), and radial diffusivity (RD) were calculated for each segmented hypothalamic nucleus, serving as distinct regions of interest. No significant differences in MD or RD were observed between males and females, nor were there lateralization effects within the same sex for these metrics. Conversely, FA—an indicator of orientation-dependent water diffusion that reflects microstructural properties such as axon diameter, degree of myelination, fiber density, and fiber collimation—displayed notable sex differences (Figure 2c). Specifically, the AVPV exhibited higher FA values in females compared to males (0.139 ± 0.008 in females vs 0.114 ± 0.005 in males, respectively; Mann-Whitney test: U = 5, p = 0.0111, n = 6, 7). This aligns with existing literature in the rat indicating that the AVPV contains a higher density of fibers in females than males ^77^. Similar sex-differences were found in the MPN, with higher FA values in females than in males (0.1107 ± 0.004 in females vs. 0.088 ± 0.002 in males; Mann-Whitney test: U = 0, p = 0.0006, n = 7, 6) (Figure 2d). These differences may result from structural heterogeneity within the nucleus that varies with sex, as suggested by findings in rats^78^. Sex-related differences were also observed in the FA of the AHA (0.105 ± 0.004 in females vs. 0.095 ± 0.002 in males; Mann-Whitney test: U = 7, p = 0.0262, n = 7, 6 for the AHA) (Figure 2d). Furthermore, FA revealed a lateralization of the LPO in both sexes, with a significantly higher FA on the right side compared to the left (0.161 ± 0.008 for the left side and 0.194 ± 0.008 for the right; Mann-Whitney test: U = 5, p = 0.011, n = 6 for females; 0.144 ± 0.01 for the left side and 0.182 ± 0.006 for the right; Mann-Whitney test: U = 4, p = 0.007, n = 6 for males) (Figure 2e). In contrast, the lateralization of certain nuclei was sex-specific. The FA of the PMv was significantly higher on the right than on the left for females (0.11 ± 0.006 for the left side and 0.132 ± 0.004 for the right; Mann-Whitney test: U = 4, p = 0.007, n = 6) but not for males (0.1321 ± 0.005 for the left side and 0.1236 ± 0.009 for the right; Mann-Whitney test: U = 16, p = 0.3176, n = 7) (Figure 2d). Conversely, the FA of the MPN and MPOA was higher on the right than on the left side for males (0.083 ± 0.002 for the left side and 0.094 ± 0.002 for the right; Mann-Whitney test: U = 2, p = 0.002, n = 6 for the MPN; 0.125 ± 0.005 for the left side and 0.147 ± 0.004 for the right; Mann-Whitney test: U = 3, p = 0.004, n = 6 for the MPOA) but not for females (0.1078 ± 0.004 for the left side and 0.1136 ± 0.004 for the right; Mann-Whitney test: U = 16, p = 0.3176, n = 7 for the MPN; 0.1434 ± 0.01 for the left side and 0.1654 ± 0.01 for the right; Mann-Whitney test: U = 10, p = 0.0728, n = 7 for the MPOA) (Figure 2d).

#### Insights into the Structural Connectivity of the Mouse Hypothalamus

Next, we used diffusion-weighed MRI tractography to reconstruct the hypothalamic connectome. Specifically, we acquired *ex vivo* diffusion-weighted data with high spatial (100 μm isotropic) and angular resolutions (up to 90 diffusion-encoding directions) and used a probabilistic tractography algorithm^79^ to compute whole-brain tractograms. These were then co-registered with the hypothalamus-enriched atlas, allowing us to reconstruct structural connections between hypothalamic nuclei themselves, arranged along specific bundles^15^, and between the hypothalamus and the rest of the brain (Figure 3a-e, Figure 4_-left panels_ and Figure S1).

**Figure 3.**
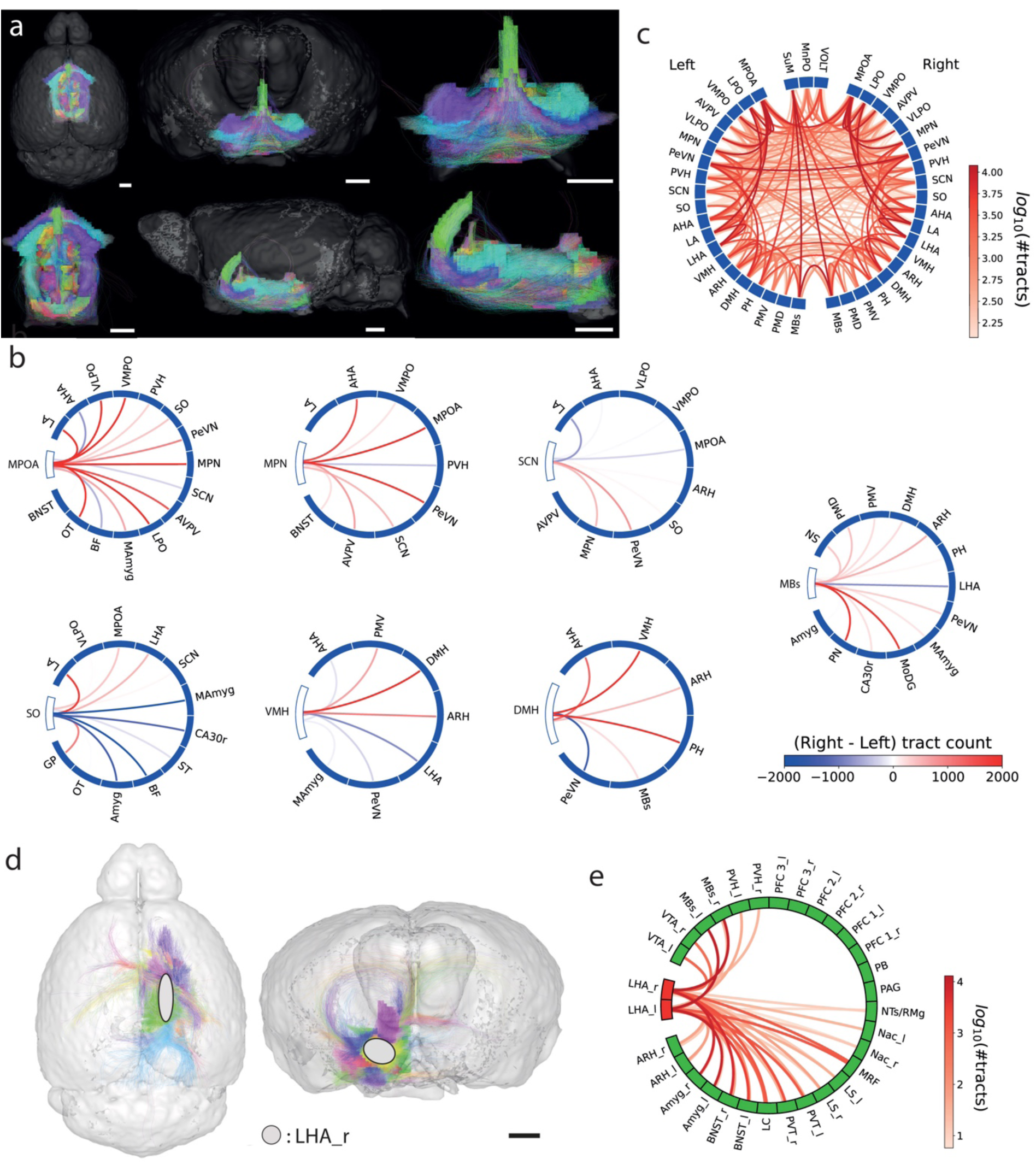
Structural Connectivity of Mouse Hypothalamic Nuclei. (**a**) Illustration of structural connectivity between hypothalamic nuclei in horizontal (left), sagittal (top right) and coronal (bottom right) planes. Streamlines are shown without applying a threshold to the number of fibers. Colors vary according to the direction in which the fibers run. (**b**) Mean hemispheric difference in the number of diffusion-derived fiber tracts (Right – Left; n = 13 mice) for the DMH, VMH, MPN and MBs found to have structural lateralization indices significantly different from zero (p < 0.005). Red lines indicate stronger right-hemisphere connectivity, blue lines indicate stronger left-hemisphere connectivity. (**c**) Structural connectivity among hypothalamic nuclei. (**d**) Illustration of structural connectivity between the LHA (ROI shown by the white ovals) and the rest of the brain in horizontal (left panel) and sagittal (right panel) sections. Streamlines are shown without applying a threshold to the number of fibers. (**e**) Structural connectivity between the LHA and selected regions reviewed in ^81^ using tractography. No threshold was used in this representation. Abbreviations are listed in Table 1. Scale bars: 1mm.

**Figure 4.**
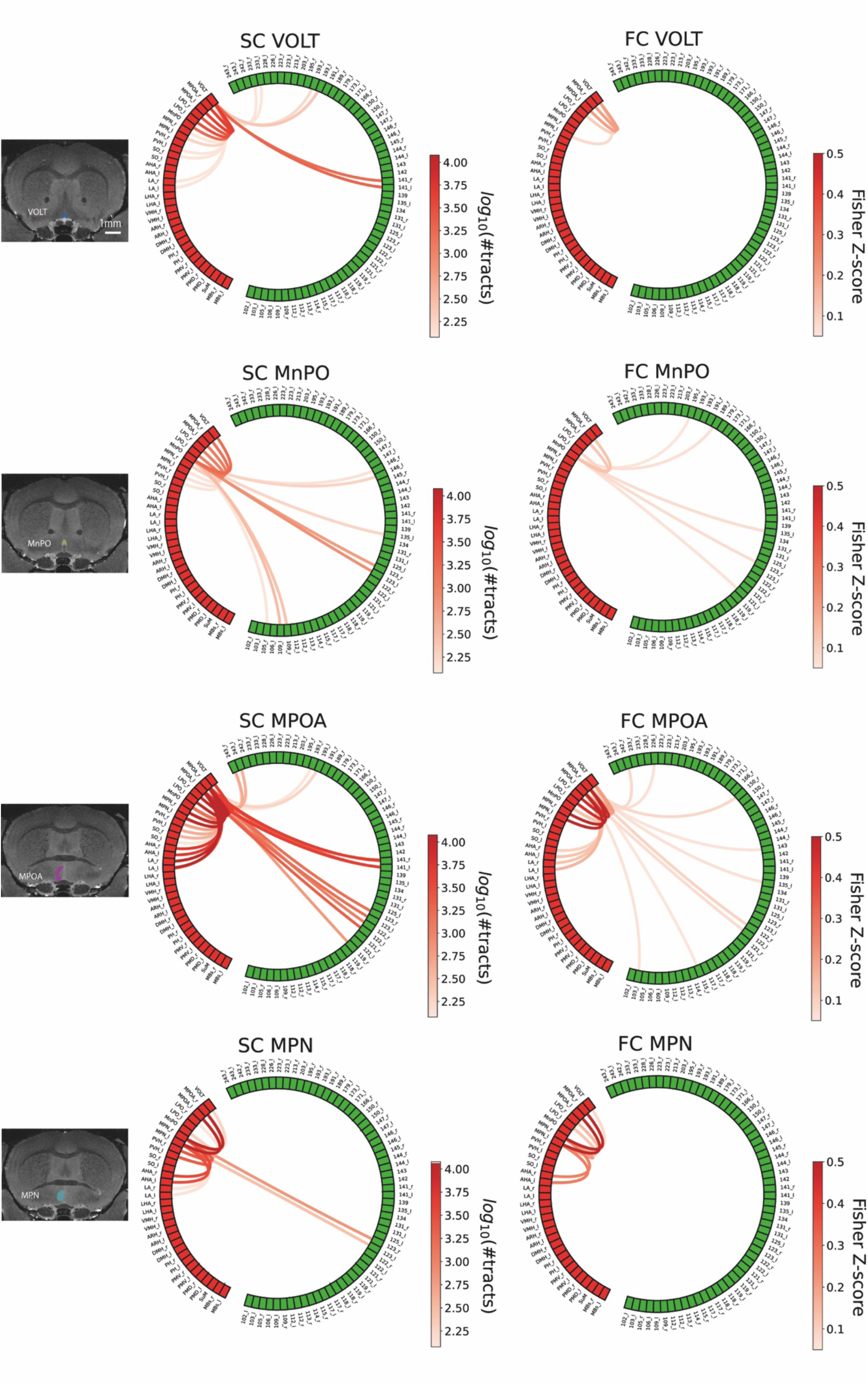
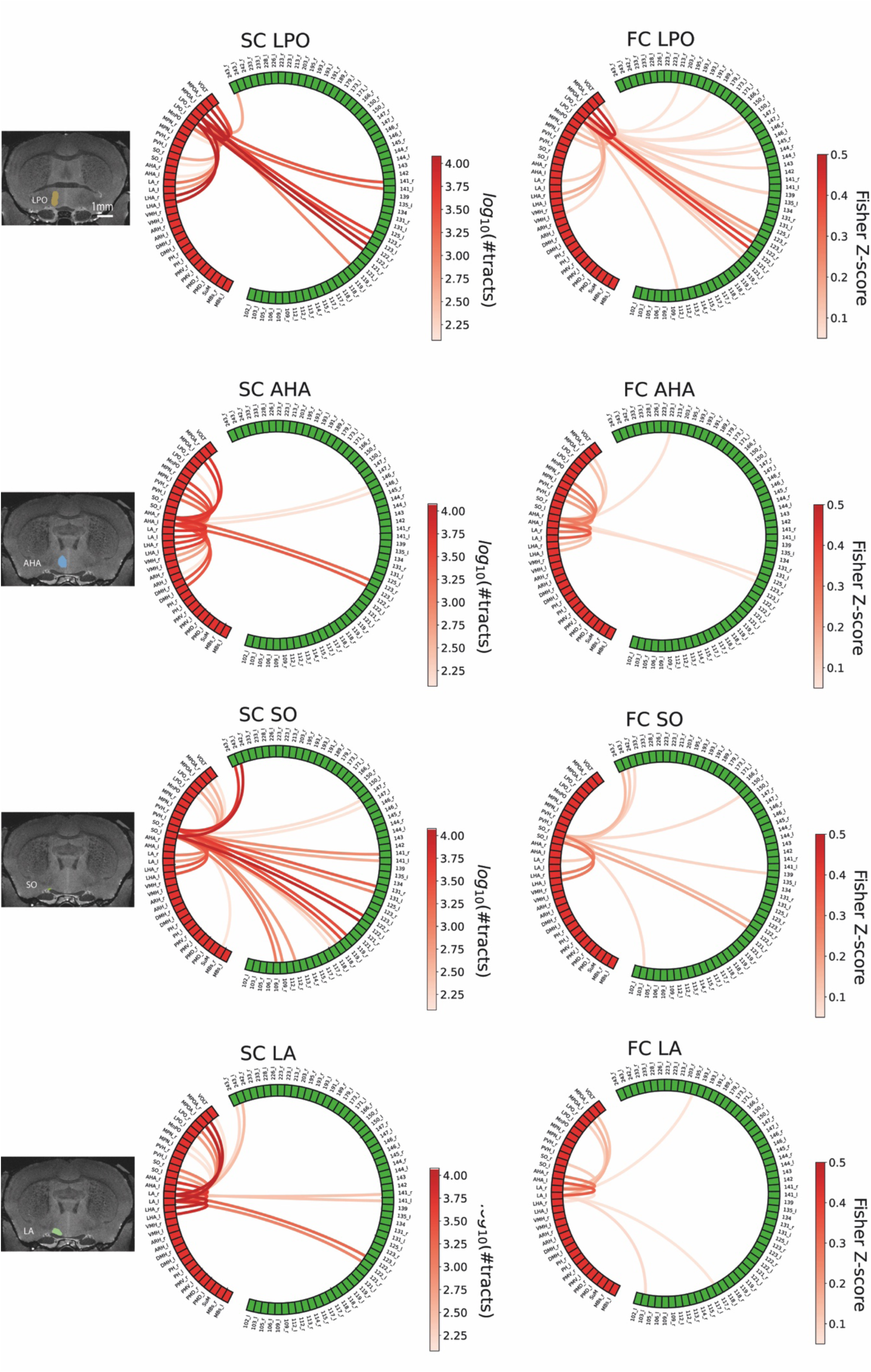
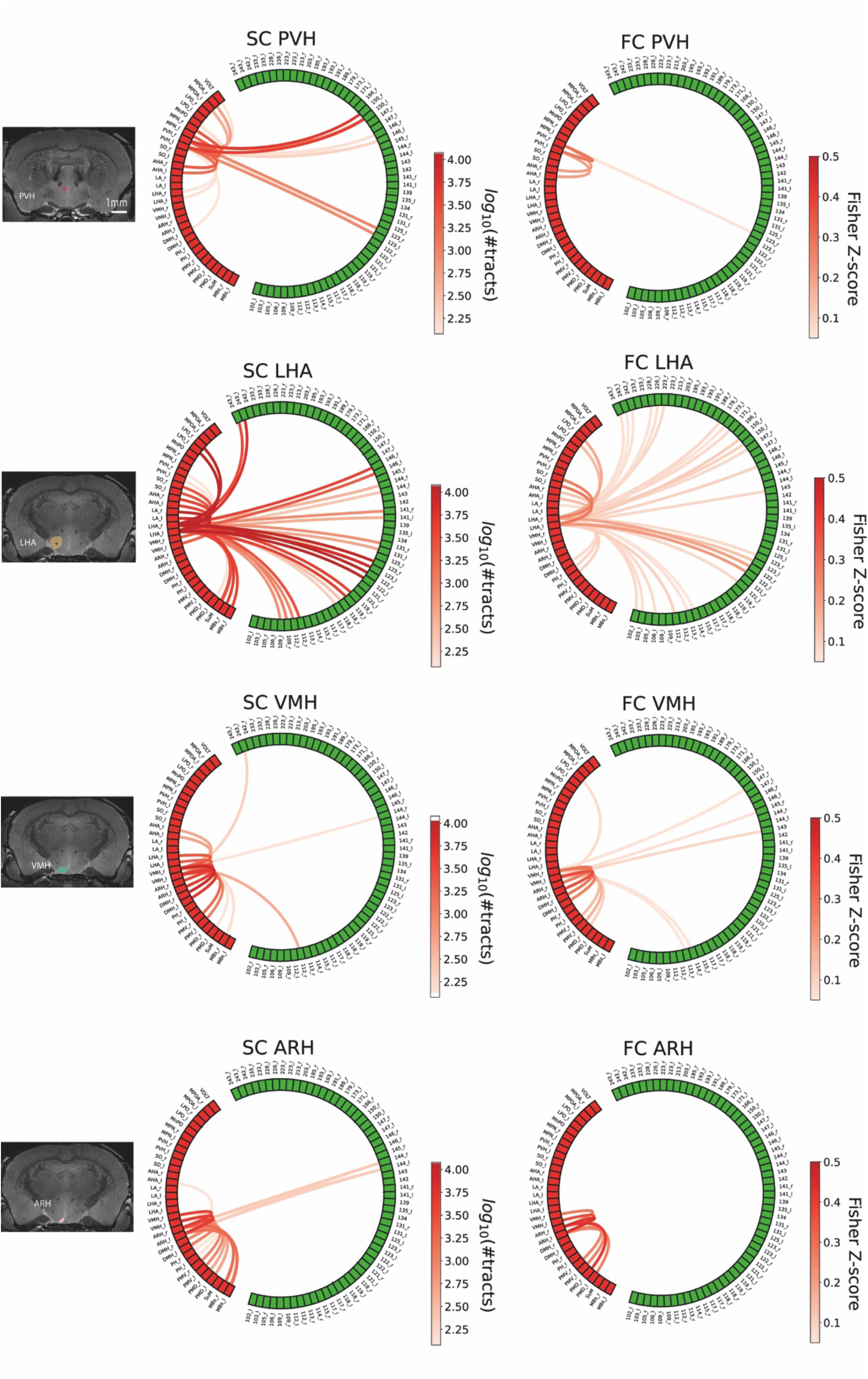
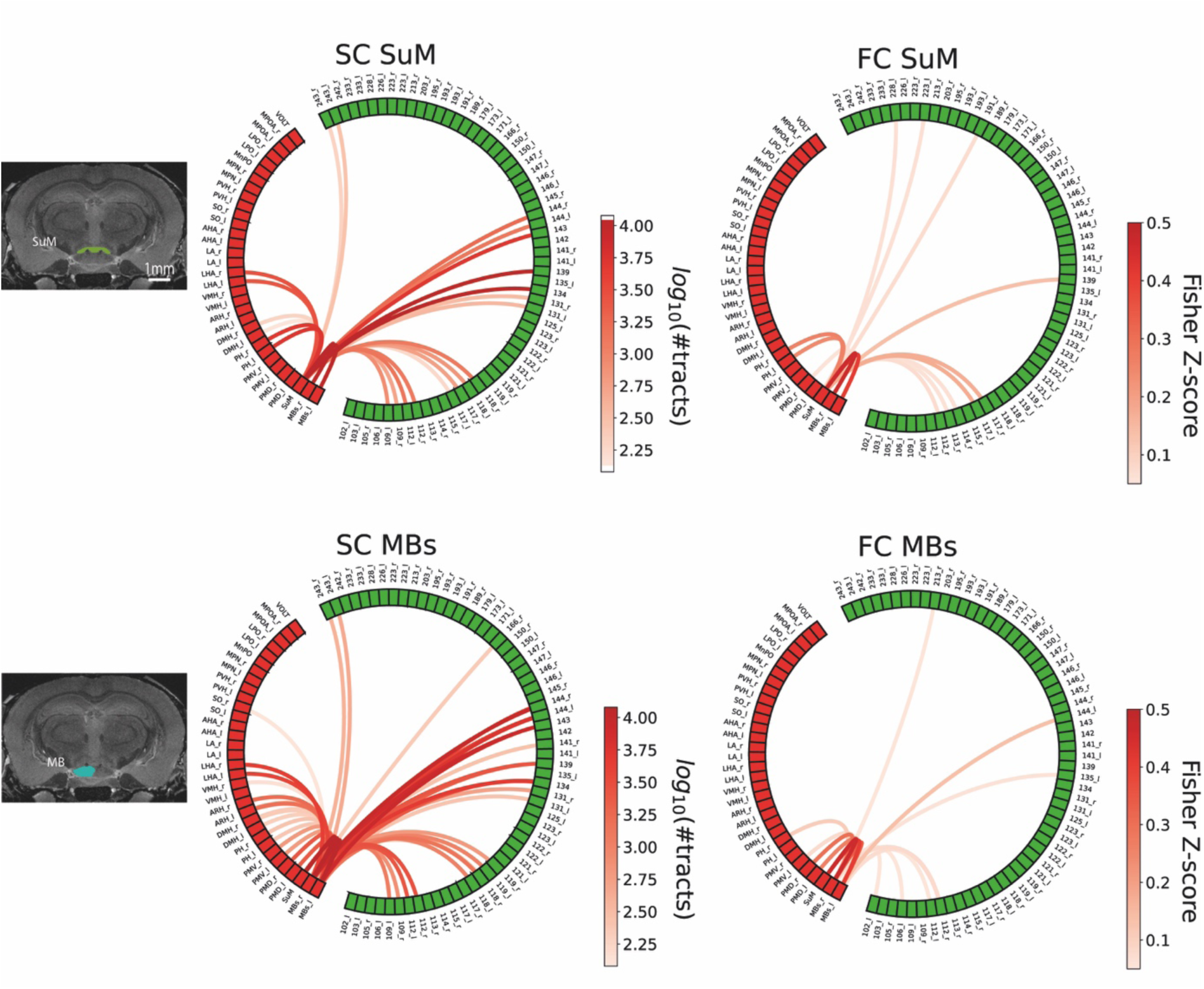
Coupling of Structural and Functional Connectivity in the Hypothalamus. Structural and functional connectivity profiles of the 14 hypothalamic nuclei for which both structural connectivity (SC) and functional connectivity (FC) could be assess. Major connections are consistently observed in both structural and functional domains with a weighted Dice score > 0.7. Abbreviations are listed in Table 1.

No significant sex differences were observed in hypothalamic structural connectivity, based on both Python-based analysis (SciPy’s Mann-Whitney U test false discovery rate (FDR)-corrected^80^, p>0.05 for all tests) and Network Based Statistics (NBS). In terms of lateralization, among the 20 hypothalamic nuclei considered, seven displayed structural lateralization indices (*sLi)* significantly different from zero (p < 0.05). These hypothalamic regions showed greater structural connectivity in the right hemisphere than in the left (Figure 3b).

A complementary, network-level view of structural connectivity between hypothalamic nuclei is shown in Figure 3c.

We compared our data with existing studies on the long-range connectivity of the lateral hypothalamic area (LHA) using standard tracing methods (reviewed in ^81^). Because some identified regions were not individually segmented in our atlas, we adapted our analysis by using, when necessary, larger anatomical regions that encompassed them or by clustering several small regions. For example, cingulate cortex areas were used in place of the prefrontal cortex while the dorsal raphe nucleus, which is not segmented in the DSURQE MRI atlas, had to be excluded. Our dataset confirmed major LHA connections documented in the literature, including projections from the bed nucleus of the stria terminalis (BNST), which descend through the medial forebrain bundle^82, 83^ and connect the LHA with the amygdala, the paraventricular nucleus of the hypothalamus (PVH), the paraventricular nucleus of the thalamus (PVT), the ventral tegmental area (VTA), the midbrain reticular formation (MRF) and finally, the nucleus tractus solarius (NTS) in the brainstem, among others (Figure 3d, 3e). Additionally, in line with known projections from the LHA ^84^, we observed structural connectivity with the lateral septum (LS) and several areas within the hypothalamus, including the LPO, the AHA, the VMH and the SuM, as well as weaker connectivity to the dorsal premammillary nucleus (PMd) (Figure 3d, Figure 4_-left panel_). However, we did not detect projections to the prefrontal cortex, parabrachial nucleus or the DMH (Figure 4_-left panel_). The absence of signal from these pathways could be due to factors such as species differences, complex fiber geometry or limitations to the sensitivity of diffusion tractography, which may not detect the presence of a very low number of fibers.

Unexpectedly, our dataset highlights a strong structural connectivity between the ARH and the VMH (Figure 4_-left panel_) which is in contrast to previous observation of the paucity of the projections between the VMH and the ARH ^85^ ^86^. This connectivity may correspond to the cell-poor region that separates the VMH from the dorsal surface of the ARH, documented to innervate the ARH ^86^. Additionally, the extension of ARH dendrites into the VMH, as supported by early Golgi staining studies ^87^ may contribute to this structural relationship between the two hypothalamic nuclei. The structural connectivity between the ARH and the VMH may also partly be explained by the extension of the distal processes of tanycytes, which constitute a substantial portion of total tissue volume medially^87^. These processes have been shown to extend laterally through the parenchyma ^87^, potentially forming a bridge from the most dorsal portion of the ARH to the VMH (see Figure 3A in ^88^).

#### Insights into the Functional Connectivity of the Mouse Hypothalamus

High-magnetic-field functional MRI (fMRI) offers improved contrast and a better correlation with underlying neuronal activity due to the increased contribution of small blood vessels to the blood-oxygen-level dependent (BOLD) signal^89^. The independent component analysis (ICA)^90^ revealed previously reported networks^91, 92^ (Figure S2) as well as hitherto undescribed networks extending into hypothalamic regions (Figure 5a). However, the same methodology that provided better contrast also led to greater susceptibility-related artifacts in regions near air–tissue interfaces. To minimize these artifacts, we applied dynamic shimming and TOPUP correction^93, 94^. Despite these efforts, some hypothalamic nuclei — including the PeVN, SCN, VLPO, VMPO and AVPV, which were identifiable in the high-resolution anatomical images — were not distinguishable or significantly cropped in the resting-state fMRI images (rs-fMRI) and therefore excluded from the functional connectivity analysis (Figure 5b). The VMH was partially altered by susceptibility artifacts but still exploitable. Pearson’s correlation coefficients were first calculated between each pair of regions of interest (ROIs) in the atlas and then converted to Fisher z-scores. No significant differences in functional connectivity were found between male and female groups, either within the hypothalamus or between the hypothalamus and the rest of the brain, using FDR-corrected Mann-Whitney U tests (p>0.05 for all tests). No sex differences were observed in the resting-state networks identified by group ICA^90^ using FSL’s dual regression^95^. Therefore, data from male and female subjects were pooled and connectivity matrices were constructed from Fisher-transformed correlation values, retaining only significant connections (Mann-Whitney U, FDR-corrected, p<0.01) (Figure 5c, 5d). Functional connectivity was then mapped both between hypothalamic nuclei themselves (Figure 5c) and between hypothalamic nuclei and whole-brain gray matter (Figure 5d).

**Figure 5.**
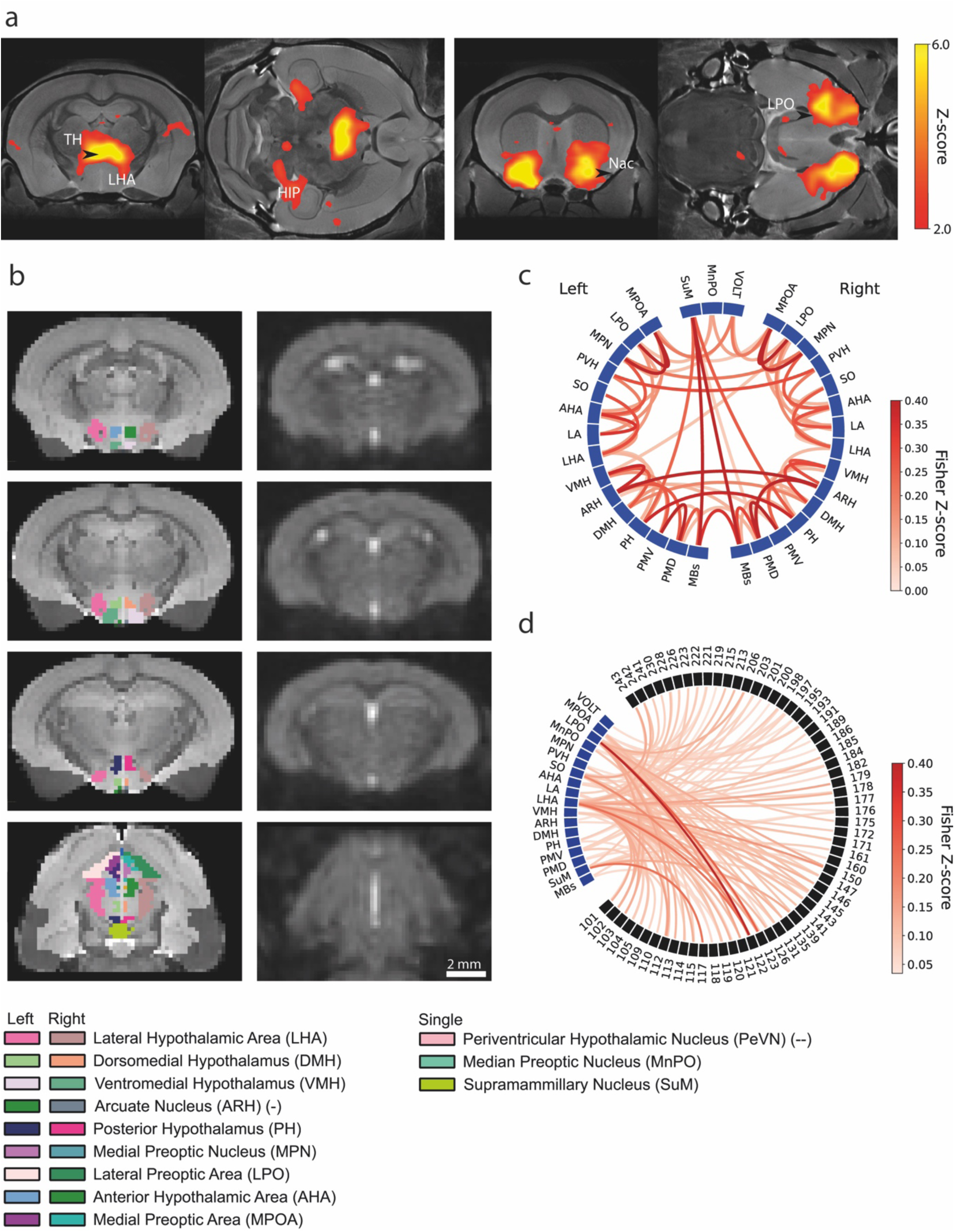
Functional Connectivity of Mouse Hypothalamic Nuclei. (**a**) Independent component analysis highlighting functional components involving the diagonal bands of Broca and the lateral preoptic nucleus, among other regions, overlaid on the anatomical template. (**b**) Coronal (top three rows) and horizontal (bottom row) sections of the mouse brain extracted from anatomical (left) and functional (right) acquisitions. Brain areas successfully captured in fMRI are highlighted in lighter gray in the left column. Regions marked with a (-) are partially altered by susceptibility artifacts but still exploitable, whereas regions marked with a (--) are fully lost due to these artifacts or undersampling to fMRI resolution. (**c**) Statistically significant functional correlations among hypothalamic nuclei, using Mann-Whitney U test on Fisher-Z transformed Pearson correlation coefficients (p<0.01, FDR corrected, n= 13). (**d**) Statistically significant functional correlations between hypothalamic nuclei and other regions of the brain segmented in HypoAtlas, analyzed using the same statistical approach as in (c). Abbreviations are listed in Table 1.

#### Coupling of Structural and Functional Connectivity in the Hypothalamus

Connectivity information from diffusion tractography and rs-fMRI was used to assess structure–function coupling in the hypothalamus using the weighted-DICE–based metric, computed on group-averaged connectivity matrices. The resulting structural connectivity (SC)–functional connectivity (FC) coupling exceeded 0.7 (out of 1) for all hypothalamic nuclei studied, except for the MnPO and PVH, for which the weighted Dice score remained above 0.5. The corresponding group-averaged structural and functional connectograms are shown in Figure 4 and Supplementary Figures 1. Importantly, several dominant connectivity patterns were consistently observed across both modalities. Moreover, consistent with established neuroanatomical findings demonstrating the subiculum’s connection to the hypothalamus through the corticohypothalamic tract^96^, but not detected by our structural connectivity paradigm, functional connectivity was identified between the subiculum and the MPOA (Figure 4). Similarly, functional connectivity was shown between the LHA and the DMH, which are known to be structurally connected^84, 97^ but not detected by our structural connectivity paradigm (Figure 4). Notably, the substantial structural connectivity observed between the VMH and the ARH was complemented by a strong functional connectivity (Figure 4). This is in line with previous studies that used advanced laser scanning photostimulation techniques to reveal that ARH neurons—POMC neurons in particular—were synaptically activated by VMH neurons ^98^. Additionally, it has recently been shown that the chemogenetic manipulation of tanycytes can induce neuronal activation in the ARH parenchyma^99^, underscoring the complex interplay within this neural network^100^. It is worth noting that tanycytic processes not only encircle and occasionally terminate on blood vessels, regulating their permeability^101, 102^, and hence vascular dynamics. This raises the intriguing prospect that tanycytes could dynamically modulate the BOLD signal in the ARH-VMH region of the tuberal hypothalamus.

#### Sex Differences in Hypothalamic Metabolic Profiles Revealed by ¹H-MR Spectroscopy

For spectroscopic measurements, we positioned our voxel to include both sides of the hypothalamus, with a volume of interest (VOI) of 2.1 mm x 1.3 mm x 2.4 mm corresponding to 6.5µL as shown in Figure 6a. At 17.2 T field strength, we consistently obtained high quality spectra (Figure 6b), with metabolite concentrations in the hypothalamus often higher than those seen at 14.1 T ^103^ and, notably, serine, a metabolite previously not reliably detected by MRS, was clearly resolved in every animal’s ^1^H-Nuclear magnetic resonance (^1^H-NMR) spectrum (Figure 6b).

**Figure 6.**
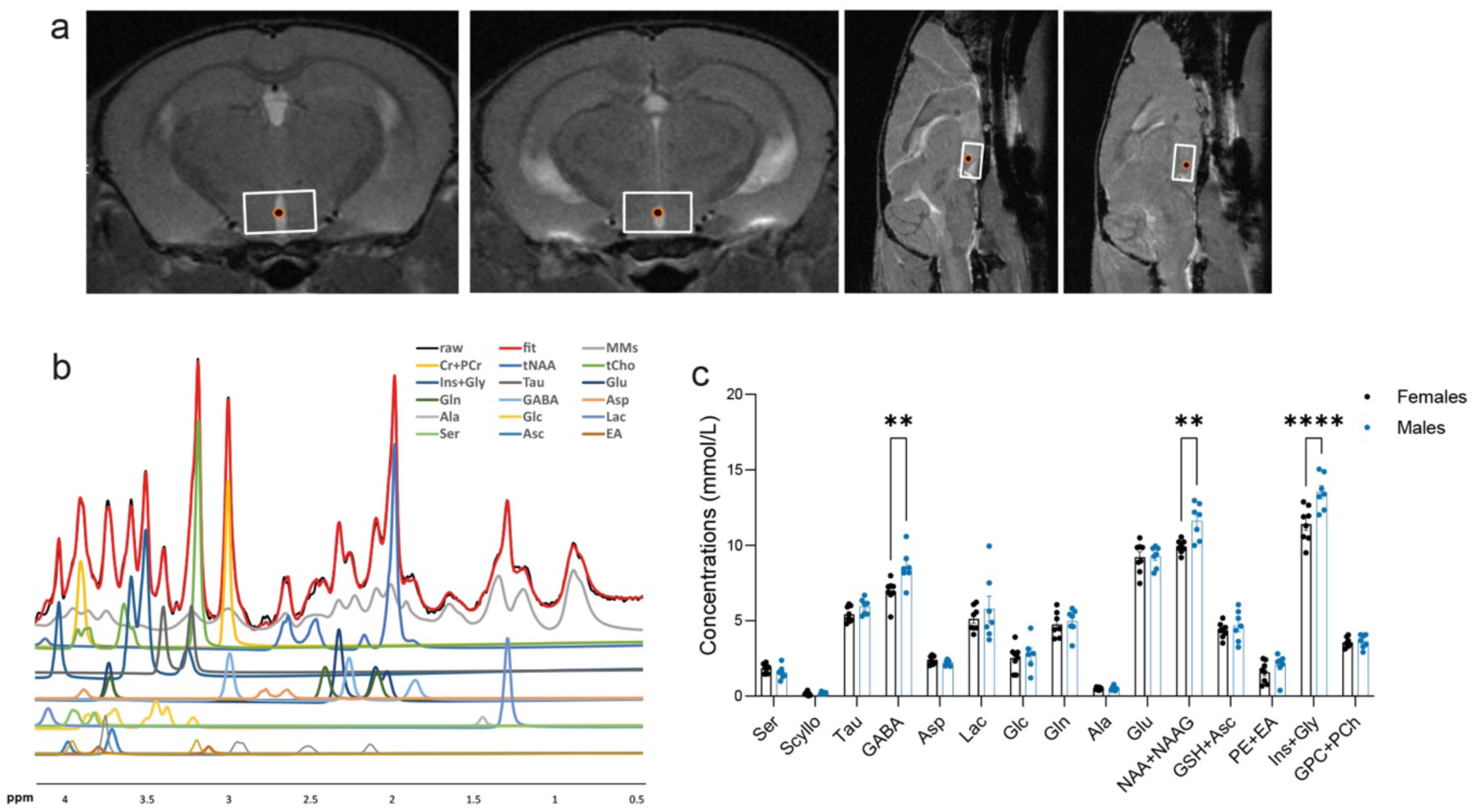
Localized ^1^H NMR spectra of mouse hypothalamus at 17.2. **T**. (**a**) Voxel positions in coronal and sagittal orientations, covering the tuberal, posterior, medial and lateral parts of the hypothalamus (voxel size: 2.1 x 1.3 x 2.4 mm^3^, 6.5µl). The volume of interest (VOI) is marked in white. (**b**) Localized ^1^H NMR spectra obtained from the VOI. Each metabolite or sum of metabolites is indicated by a color and the fit of all metabolites is represented in red. (**c**) Neurochemical profiles of the mouse hypothalamus in females (black bars) and males (blues bars). All measured metabolites are shown, with significant sex differences indicated (Bonferroni multiple comparisons test, **p = 0.028 for GABA, **p = 0.032 for NAA+NAAG and ****p = <0.0001 for Ins + Gly). Metabolites concentrations were normalized to Cr+PCr. Abbreviations: Ser: serine, Gly: glycine, Asc: ascorbate, Scyllo: scyllo-inositol, GSH: glutathione, PCh: phos-phocholine, GPC : glycerophoshocholine, Tau: Taurine, Ins: myoinositol , Glu: glutamate, NAA : N-actetyl-aspartate, NAAG: N-acetyl-aspartyl-glutamate, Gln: glutamine, GABA : γ-amino-butyric acid, Asp: aspartate, Cr: creatine, PCr phosphocreatine, Lac: lactate, Glc: glucose, Ala: alanine, EA: ethanolamine, PE: phosphoethanolamine.

Our results indicate that the concentrations of certain metabolites are higher in the hypothalamus of males compared to females in the diestrous stage of their estrous cycle (when circulating estrogens are at their lowest), including γ-amino-butyric acid (GABA) (8.4 ± 0.4 mmol/L in males vs 6.9 ± 0.3 mmol/L in females, Bonferroni’s multiple comparisons test, t = 3.846, p = 0.024, n = 7, 8), the sum of N-acetyl-aspartate (NAA) and N-acetyl-aspartyl-glutamate (NAAG), a marker of neuronal activity and viability^104^ (11.3 ± 0.5 µmol/g for males vs. 9.9 ± 0.2 mmol/L in females, Bonferroni’s multiple comparisons test, t = 4.048, p = 0.011, n = 7, 8) and the sum of myoinositol, a marker of glial size and activity^105, 106^, and glycine, a gliotransmitter acting as a co-agonist of extrasynaptic N-methyl-D-aspartate (NMDA) receptors^107^ (13.0 ± 0.7 µmol/g for males vs 11.4 ± 0.4 mmol/L for females, Bonferroni’s multiple comparisons test, t = 5.045, p = <0.0001, n = 7, 8) (Figure 6c). No differences were observed in other metabolites examined, including glucose, macromolecules and lipids.

## Discussion

In this study, we employed ultra-high-field 17.2 T MRI to comprehensively investigate the structure, connectivity, function and neurochemistry of the mouse hypothalamus. By combining high-resolution anatomical imaging, *ex vivo* diffusion MRI and tractography, *in vivo* resting-state fMRI, and *in vivo* ¹H MRS, we characterized this small yet highly functionally diverse brain region with unprecedented detail. Diffusion and tractography analyses were enabled by expert manual segmentation of the hypothalamus and the development of a novel atlas of mouse hypothalamic nuclei. Importantly, we have made this segmentation (HypoAtlas) publicly available through the EBRAINS digital research infrastructure, developed as part of the Human Brain Project ^108, 109^.

The HypoAtlas captures nuclei that are unresolved in existing MRI atlases, providing a more precise and comprehensive anatomical framework for future structural and functional analyses of the hypothalamus. Using *ex vivo* diffusion-weighted imaging and tractography, we mapped both intrahypothalamic pathways and connections to other brain regions. The detailed connectivity maps presented here will facilitate future circuit-level investigations, eventually enabling the discovery of previously unrecognized structural connections both within the hypothalamus and between hypothalamic nuclei and extrahypothalamic structures, without preconceived assumptions.

*In vivo* resting-state fMRI combined with ICA also revealed functional components involving hypothalamic nuclei. Some of these components have not been previously reported in the mouse literature, suggesting that the increased spatial resolution and BOLD contrast-to-noise ratio at 17.2 T enables detection of subtle, deep-brain resting-state networks. This underscores the value of high-field fMRI for exploring the functional organization of deep brain structures, which are often underrepresented in functional connectivity studies.

We also acquired *in vivo* ^1^H-NMR spectra over the entire hypothalamus, yielding a neurochemical profile that aligns well with previous ultra-high-field studies, particularly the work by Lei *et al*.^103^. A previous study using ultra-high-field 14.1 T MRI^103^ successfully characterized mouse hypothalamic metabolites but without comparisons between the sexes. More recently, Tkac *et al*.^110^ have studied the profile of mouse hypothalamic metabolites according to sex at 9.4 T but found no significant differences. In our study, the use of ultra-high field MRI not only increased sensitivity and spectral resolution, but also allowed for unprecedented systematic detection of low-concentration metabolites such as serine and glycine (Figure 6b), validating the feasibility of *in vivo* hypothalamic MRS at ultra-high field.

We also observed sexual dimorphism in both anatomy and diffusion metrics within the hypothalamus; importantly, the ultra-high-field MRI resolution achieved here enabled detection of these differences at the level of individual hypothalamic nuclei, which, to our knowledge, has not previously been demonstrated *in vivo*. Our observations further support the need for sex-aware experimental design in preclinical neuroscience. Finally, weighted Dice values were generally highly similar between structural and functional connectivity networks, with most hypothalamic nuclei showing scores above 0.7 and a few regions (MnPO and PVH) above 0.5, indicating good overall agreement between diffusion-based and rs-fMRI–derived connectivity patterns at the group level.

There are several limitations to consider. Despite dynamic shimming and post-acquisition distortion correction, the use of echo-planar imaging (EPI) for fMRI resulted in signal dropout in ventral brain areas, including parts of the amygdala and lower hypothalamus, due to susceptibility artifacts at air–tissue interfaces. Additionally, while our ¹H-MRS data provide valuable insight into hypothalamic neurochemistry, the spectroscopy voxel covered the entire hypothalamus and thus lacked subregional specificity. Moreover, because structural and functional connectivity were visualized in different groups of mice, their coupling could only be assessed at the group level, rather than at the individual level. A further limitation is that the similarity in Dice scores was computed after applying a streamline-count threshold to the diffusion tractography data, which may influence the estimated degree of structure–function coupling. This thresholding step, although applied for robustness, may bias coupling estimates by excluding weaker or less consistently reconstructed connections.

Taken together, our results reveal the advantages of ultra-high-field MRI for the characterization of the mouse hypothalamus across structural, functional and neurochemical dimensions. The refined anatomical atlas, connectivity maps, resting-state networks and spectroscopy data reported here provide a valuable resource for future studies of hypothalamic circuits and functions. Our findings also underscore the critical importance of incorporating sex as a fundamental biological variable in hypothalamic research, and demonstrate that these sex-dependent features can be interrogated in vivo using ultra-high-field 17.2 T MRI. Additionally, they open new avenues for investigating the role of the hypothalamus in adaptive physiology, neurodevelopmental and age-related brain disorders, including those arising from brain-body miscommunication, particularly by making it possible to conduct longitudinal structural and functional studies in the same mice.

## Materials and Methods

All animal procedures were approved by the Committee for Ethics in Animal Experimentation, Commissariat à l’Energie Atomique et aux Énergies Alternatives (CEA), and by the French Ministry for Higher Education and Research under reference #41874-2023032112005163 v1, and were conducted in strict accordance with the recommendations and guidelines of the European Union (Directive 2010/63/EU) and the French National Committee (Décret 2013–118). Mice (C57BL/N6, Janvier Labs, Le Genest Saint Isle, France) were fed *ad libitum* and housed under a 12-hour light-dark cycle at 22°C and 50% humidity. A total of 43 mice (22 males and 21 females) from 2 to 4 months of age were included in this study. All acquisitions were performed using a 17.2 Tesla MRI scanner (Bruker Biospin GmbH, Ettlingen, Germany) equipped with a gradient set of maximum strength of 1 T/m. A transmit/receive quadrature birdcage coil with a 2.4 cm inner diameter (Rapid Biomedical GmbH, Rimpar, Germany) was used for both *in vivo* and *ex vivo* acquisitions.

### Ex vivo MRI

#### Tissue preparation

After transcardiac perfusion with 4% paraformaldehyde in phosphate buffered saline (PBS) 0.1M pH 7.4 and 0.4 % gadolinium (Gd-DOTA, Clariscan®), brains (within the skull) were collected and post-fixed by immersion in 4% paraformaldehyde + Gd-DOTA for 24 hours. After fixation, the brains were placed in PBS 0.1M, pH 7.4 + 0.4 % Gd-DOTA for storage. For imaging, the samples were placed in a 14 ml Falcon tube filled with Fluorinert (FC40, Sigma-Aldrich, L’Isle d’Abeau Chesnes, France).

#### Anatomical and diffusion MRI acquisitions

High resolution anatomical images, 40 µm isotropic, were acquired with a 3D T_2_-weighted Rapid Acquisition with Refocused Echoes (RARE) sequence with the following acquisition parameters: Repetition time / Time to Echo (TR/TE) 1500/12 ms, RARE factor 4, NA 2. For structural characterization, diffusion weighted images were acquired using a 3D diffusion-weighted Pulsed-Gradient Spin Echo EPI sequence with the following parameters: 14 segments, TR/TE 250/26 ms, diffusion gradient duration/spacing δ/Δ 5.0/13.3 ms, b-values 2/4/6/8/12/14/16/19 ms/µm² and 42/42/42/90/42/42/42/42 diffusion directions, 30 b=0 ms/µm² non-diffusion weighted volumes, 100 µm isotropic resolution, total acquisition time 57 hours.

### In vivo MRI

#### Animal preparation

For fMRI imaging sessions, mice were sedated with a continuous perfusion of dexmedetomidine as follows: induction of anesthesia was done with 3% isoflurane and then decreased gradually down to 0% within 30 min; dexmedetomidine was administered as a bolus (0.025 mg/kg subcutaneous) at the beginning of the session and a subcutaneous perfusion (0.1 mg/kg/h) for the entire duration of the session. During spectroscopy imaging sessions, mice were anesthetized with isoflurane (1.2 – 1.8%). Mice were placed in a cradle with an integrated water chamber^111^ and the head was immobilized using a bite bar and ear pins. The body temperature of the animals was maintained in the range of 36–37.5°C by connecting the cradle to a heated circulating bath (TC120, GRANT Instruments, Amsterdam, The Netherlands). Temperature and respiration rate were continuously monitored using a small-animal monitoring system (Model 1040, SA Instruments, Stony Brook, USA). For all acquisitions, the mice were freely breathing an air–oxygen gas mixture (33 % O_2_).

#### fMRI acquisitions

A RARE sequence was used to acquire anatomical images for 5 echo times equally spaced between 6 and 55 ms. Other relevant sequence parameters are as follows: TR 3500 ms, field of view (FOV) 1.5 x 1.5 cm, in-plane resolution 100 μm^2^, slice thickness 300 μm, 41 coronal slices, RARE factor 2. The five scans were added together to produce a reference anatomical image. A B_0_ field map was acquired prior to the RARE acquisition in order to perform shimming over the area of interest. Two BOLD rs-fMRI time series were acquired using a single shot Gradient Echo Echoplanar (GE-EPI) sequence with the following parameters: TR/TE 1500/11 ms, flip angle 55°, FOV 1.28 × 1.28 cm, in-plane resolution 173 µm^2^, slice thickness 350 μm, 25 coronal slices, 400 repetitions. In addition, a time series consisting of 40 repetitions was acquired with the phase encoding direction reversed to correct for magnetic susceptibility distortions. For the rs-fMRI acquisitions, the B_0_ field homogeneity optimization was performed using dynamic slice-by-slice shimming.

#### MRS acquisitions

Localized ^1^H-NMR spectra were acquired from a 6.5 µL volume of interest (VOI) centered on the hypothalamus (Figure 6) using a LASER (Localization by Adiabatic SElective Refocusing^112^ sequence) (TE/TR =25/3500 ms, 32 averages, 16 repetitions, spectral bandwidth 8 kHz, 4096 complex points) consisting of a 1 ms non-selective excitation pulse (10 kHz bandwidth) followed by 3 pairs of slice-selective AFP HS4 inversion pulses (2 ms, 20 kHz bandwidth). The resulting chemical shift artifact was a spatial shift of 13% between water and lipids. Prior to signal acquisition, first- and second-order shims were adjusted to optimize local B_0_ field homogeneity using the standard MAPSHIM approach in Paravision leading to typical water linewidths of 14.38 +/- 2.2 Hz. Non-suppressed water spectra were acquired for water referencing and eddy current correction. Water signal was suppressed using a VAPOR (VAriable Power and Optimized Relaxations delays) module^113^.

### Data processing

#### Segmentation of hypothalamic nuclei

A template was generated from twelve T_2_-weighted anatomical images (40 µm isotropic resolution) using the antsMultivariateTemplateConstruction2 function from ANTs ^114^, with N4 bias field correction disabled (−n 0) and the drift-prevention parameter −y set to 0. For consistency with our previous work based on the DSURQE template and atlas, the resulting template was reoriented to match the DSURQE orientation using the FSL tools fslswapdim and fslorient ^115^. The reoriented template was subsequently co-registered to the DSURQE reference space using antsRegistrationSyN, incorporating the DSURQE brain mask to improve the transformation.

Using the Paxinos atlas as a guide^28^, we manually segmented each hypothalamic nucleus visible in our template. The nuclei were identified visually based on anatomical landmarks derived from histological cross-sections, with additional guidance from previously published atlases, including the DSURQE, the Allen Brain Atlas and the Franklin and Paxinos atlas. Of note, compared to the Paxinos atlas, both our template and the DSURQE atlas exhibited a slight ventral-dorsal tilt in sections encompassing the hypothalamus, such that the dorsal part of the brain appears systematically shifted anteriorly by several slices in the MRI. This indicates a distortion between the histological atlas and the actual brain structure within the skull, which, however, did not impair the accurate identification of hypothalamic nuclei. The segmentation was performed using Itksnap software^116^ (www.itksnap.org) and relied on consistent anatomical cues including the anterior commissure, medial septum, diagonal bands of Broca, the fornices and the optic tract. A numerical label and a dedicated color were assigned to each nucleus, distinguishing left and right nuclei in order to be able to perform lateralization studies later. The segmented nuclei are listed in Table 1.

#### Anatomical MRI processing

Anatomical MRI images were first skull-stripped using a rodent skull-stripping (RSS) neural network ^117^.The anatomical atlas template was then co-registered to each individual image, producing a transformation that allowed us to map the atlas label set onto each subject’s anatomical space. Given the known voxel dimensions in this space (40 × 40 × 40 µm^3^), the volume of each ROI was computed by counting the number of voxels assigned to it. To account for inter-individual differences in brain size, ROI volumes were subsequently normalized to the total brain volume of each mouse. This yielded normalized ROI volumes for all animals in the dataset. Statistical analyses were then performed using GraphPad Prism Software v.10.

#### Diffusion MRI processing

Diffusion MRI (dMRI) images were processed using an in-house Ginkgo diffusion analysis toolbox (https://framagit.org/cpoupon/gkg). Given the large number of segments used during the multishot 3D EPI acquisition, the diffusion-weighted data were deprived of any visible geometrical distortion, limiting the pre-processing steps to a simple correction for Rician noise using a non-local means-filtering algorithm^118^. The 30 non-diffusion weighted volumes were averaged into a single T_2_-weighted volume for reference, which was in turn used to compute a precise mask of the brain using a semi-automated procedure. The diffusion process was modeled for each voxel of the mask using a Diffusion Tensor Imaging (DTI)^119^ model based on the multiple-shell data^24^ which yielded quantitative maps of FA, MD (m^2^/s) and RD, (m^2^/s). Because DTI is only able to capture a single fiber population within a voxel, a higher-order analytical Q-Ball model^120^ was computed based on the multiple-shell diffusion MRI data (spherical harmonics order 8, Laplace-Beltrami regularization factor 0006) in order to better detect fiber crossings, thereby providing diffusion orientation distribution functions (ODFs) for each individual. These ODF maps were fed to a regularized probabilistic tractography algorithm^79^ (parameters: 8 seeds/voxel, 30µm forward step, 30° aperture angle, Gibb’s sampler temperature 1) yielding an individual whole-brain tractogram of several millions of fibers. An affine registration between the anatomical MRI and the T_2_-weighted b=0s/mm² volumes was computed to further bring the atlas label map to the native space of the quantitative diffusion MRI metrics, thereby enabling the computation of average FA, MD, RD and AD for each ROI of the atlas label map. Finally, a 455×455 symmetrical connectivity matrix was computed for each individual to retrieve fibers connecting pairs of gray matter ROIs.

To eliminate spurious or isolated connections, we applied a threshold to the number of tracts connecting each pair of regions. To determine this threshold, we computed the number of fibers connecting every region pair and normalized this value by the sum of the volumes of the two regions. For each mouse, we identified the maximum normalized fiber count across all region pairs and then averaged these maxima across animals. We retained only region pairs whose volume-normalized fiber counts exceeded one-thousandth of this average maximum. This threshold was used throughout, unless otherwise specified.

### Exploring structural connectivity lateralization

We examined lateralized hypothalamic regions to assess structural connectivity differences between the right and left hemispheres. Specifically, we investigated 20 hypothalamic regions (LPO, MPN, SO, PVH, LHA, PH, VMH, DMH, MBs, ARH, PMV, PMD, MPOA, AHA, LA, PeVN, AVPV, VMPO, VLPO, SCN). For each hypothalamic nucleus, we computed the total number of reconstructed streamlines between the right ROI and the right hemisphere, and the corresponding total number between the left ROI and the left hemisphere. A structural lateralization index was defined as the difference between the right and the left terms, normalized by their sum : *sLi(ROI)* = 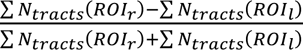. This index was computed for all hypothalamic regions and across all 13 mice scanned in diffusion. Statistical testing was then performed to assess whether these indices differed significantly from zero (Mann-Whitney U test, FDR-corrected, p<0.05).

#### fMRI processing

Raw data was denoised using the Smart Noise Reduction function in Paravision 360 (Bruker Biospin, GmbH, Ettlingen, Germany) and then converted to Nifti (Neuroimaging informatics technology initiative) format using the BrkRaw library^121^. Slice-time correction was conducted using the Analysis of Functional NeuroImages (AFNI) 3dTshift algorithm^122, 123^ followed by a deobliquing step with 3dWarp to align the oblique scans with the cardinal axes. Susceptibility-induced deformations were corrected using TOPUP^93, 94^ from FMRIB Software Library (FSL, Oxford, UK)^115^. To minimize artifacts introduced by non-linear co-registration, a preliminary harmonization step was implemented to ensure consistent spatial coverage across the atlas, functional scans and anatomical images. Specifically, functional images were co-registered to their corresponding anatomical reference image using a rigid transformation within ANTs^124^. Subject-specific brain masks were created based on the coverage of the functional scans and applied to each subject’s anatomical scan. An intersection mask was then computed across all subjects and was used to crop the atlas, retaining only regions present in every individual image. Pre-processing steps - including inhomogeneity and motion correction - were carried out using RABIES (Rodent Automated Bold Improvement of EPI Sequences)^125^. Functional images were then smoothed with a 0.3-mm full-width at half-maximum Gaussian kernel and were filtered to retain frequencies between 0.01 and 0.1 Hz. For each functional run (comprising 400 timepoints), the first and last 20 volumes were removed to avoid filtering-related edge artifacts. To remove nonneuronal fluctuations, six motion parameters and one regressor representing the average signal intensity in the cerebrospinal fluid were regressed out. White matter signals were not regressed out, given recent demonstrations of fMRI signals in white matter^126, 127^, in order to preserve potential neural contributions. Brain masks in the common space were computed for each functional scan using the RSS neural network ^117^. An intersection mask was generated by combining the functional brain masks from all subjects with the atlas mask in common space. The resulting mask included only voxels corresponding to brain tissue that were reliably acquired across the entire cohort, thereby excluding regions not consistently visible in the functional images due to susceptibility artifacts. In addition, because functional images have lower spatial resolution than the anatomical atlas, resampling the atlas into functional space can introduce resolution-related changes, particularly for small or thin structures. As a result, some ROIs may shrink, fragment or disappear. To quantify the combined impact of these effects, we examined the number of voxels retained within each ROI following resampling. Regions with insufficient voxel representation (< 6 voxels when combining the left and right ROIs for each nucleus) were excluded from further analyses (see Figure 4a for reference).

Finally, confound-corrected time series from the two runs belonging to the same subject were concatenated, yielding a single time series per subject. These datasets were used for the subsequent processing steps.

### Independent component Analysis

ICA was performed on male and female subjects separately as well as on all subjects pooled together using Melodic in FSL^90^. A dual regression analysis was used to examine between-group differences in ICA networks^95^.

### Functional connectivity

Mean time series for all ROIs in the atlas were calculated for each subject using time courses extracted from the RABIES pipeline. Functional Connectivity (FC) matrices were then computed for each subject by calculating Pearson’s correlation coefficients between time courses of ROI pairs, followed by Fisher’s Z-transformation to improve normality. Only statistically significant connections (FDR-corrected p<0.01) were retained. Network-Based Statistics (NBS)^128^ and Mann-Whitney U tests were used to evaluate potential sex differences in functional connectivity in various networks, namely within the hypothalamus and between the hypothalamus and the rest of the brain. As no significant differences were observed between male and female subjects, all participants were subsequently analyzed as a single group.

#### Exploring functional connectivity lateralization

Similar to structural connectivity, we examined lateralized hypothalamic regions to assess functional connectivity differences between the right and left hemispheres. We investigated 15 hypothalamic regions (LPO, MPN, SO, PVH, LHA, PH, VMH, DMH, MBs, ARH, PMV, PMD, MPOA, AHA, LA) and designed a functional lateralization index. For each ROI, we computed the mean Fisher’s Z-transformed functional connectivity between the right ROI and the right hemisphere, and the corresponding mean value for the left ROI and left hemisphere. A functional lateralization index was then defined as the difference between the right and left mean, normalized by their overall mean: 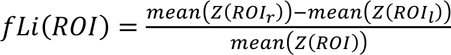. This index was computed for all 15 hypothalamic regions and across all 13 mice scanned with rs-fMRI. For the regions analyzed, none of the functional lateralization indices differed significantly from zero (Mann-Whitney U test, FDR-corrected, p>0.05 for all tests).

### Linking structural and functional brain networks (SC-FC coupling)

To quantify the SC-FC coupling, we designed a metric akin to a weighted Dice score. Functional connectivity edges were weighted by their Fisher Z-scores, while structural connectivity edges were weighted by the number of tracts connecting each pair of regions. SC–FC coupling was computed on group-averaged structural and functional connectivity matrices, obtained by averaging connectivity weights across animals within each modality. For each seed ROI, both sets of weights were normalized such that the sum of weights equaled 1. For hypothalamic nuclei with left and right hemisphere counterparts, we report the average score across the two nuclei. This weighted Dice-based metric captures the overlap between structural and functional connectivity patterns while accounting for connection strength, making it well suited for group-level SC-FC coupling analysis.

#### MRS processing

MRS data were preprocessed using Fourier transformation, zero-order phase correction, frequency shift correction and residual water signal suppression using the Hankel Lanczos singular value decomposition (HLSVD) algorithm^129^. They were then analyzed using LCModel 6.3^130^ and a basis set of simulated spectra accounting for the spectral contributions of up to 20 neurometabolites: aspartate (Asp), ascorbate (Asc), creatine (Cr), ethanolamine (EA), γ-amino-butyric acid (GABA), glucose (Glc), glutamate (Glu), glutamine (Gln), glycine (Gly), glutathione (GSH), glycero-phosphocholine (GPC), phosphocholine (PCh), myoinositol (Ins), lactate (Lac), N-acetyl-aspartate (NAA), N-acetyl-aspartyl-glutamate (NAAG), phosphocreatine (PCr), phosphoethanolamine (PE), serine (Ser), scyllo-inositol (Scyllo) and taurine (Tau). An optimized set of Gaussian basis functions was used to account for lipid and macromolecule signals^24^. Metabolite concentrations were derived from the total creatine concentration, considered as an internal reference of concentration ([tCr]= 8 mmol.L^-1^).

## Data availability statement

The hypothalamic segmentation is available via HypoAtlas on the EBRAINS platform. All other data supporting the findings of this study are available from the corresponding authors upon reasonable request.

## Acknowledgments

We thank Melissa Glatigny for support with *in vivo* MRI experiments. This study was supported by the Agence National de la Recherche (ANR, France) ANR-24-CE16-3311 HAMBURGER to VP and LC, and the 2023 Scientific Grant Prize of the Institut de France-Fondation NRJ for Neuroscience (France) and the 2023 Scientific Grand Prize of the Fondation pour la Recherche Médicale (FRM, France) to VP. AS was a doctoral student supported by a fellowship from the Inserm and the Région Hauts-de-France. The 17.2T MRI system was supported through the Ile-de-France SESAME Large Equipment program. We are indebted to S. Rasika for editing the manuscript.

## Author contributions

A.S. participated in data acquisition by performing in vivo imaging and by preparing and acquiring the ex vivo samples, conducted statistical analyses, segmented the hypothalamic nuclei, and wrote the first draft of the manuscript.; P.L.-S. participated in data acquisition and performed image processing, image analysis, statistical analysis, and template creation; I.U. processed the tractography data; F.B. processed the spectroscopy data; C. P. design the tractography acquisition protocol; V.P. contributed to project conceptualization and supervision and provided anatomical guidance, with P.C. and P.-Y.R. assisting in that role; P.V. enabled collaboration through the provision of access to a novel enabling technology, initiating the project. L.C. contributed to project conceptualization, supervision and data acquisition and led the methodological development. All authors edited and approved the manuscript.

## Supplementary Figures

**Figure S1.**
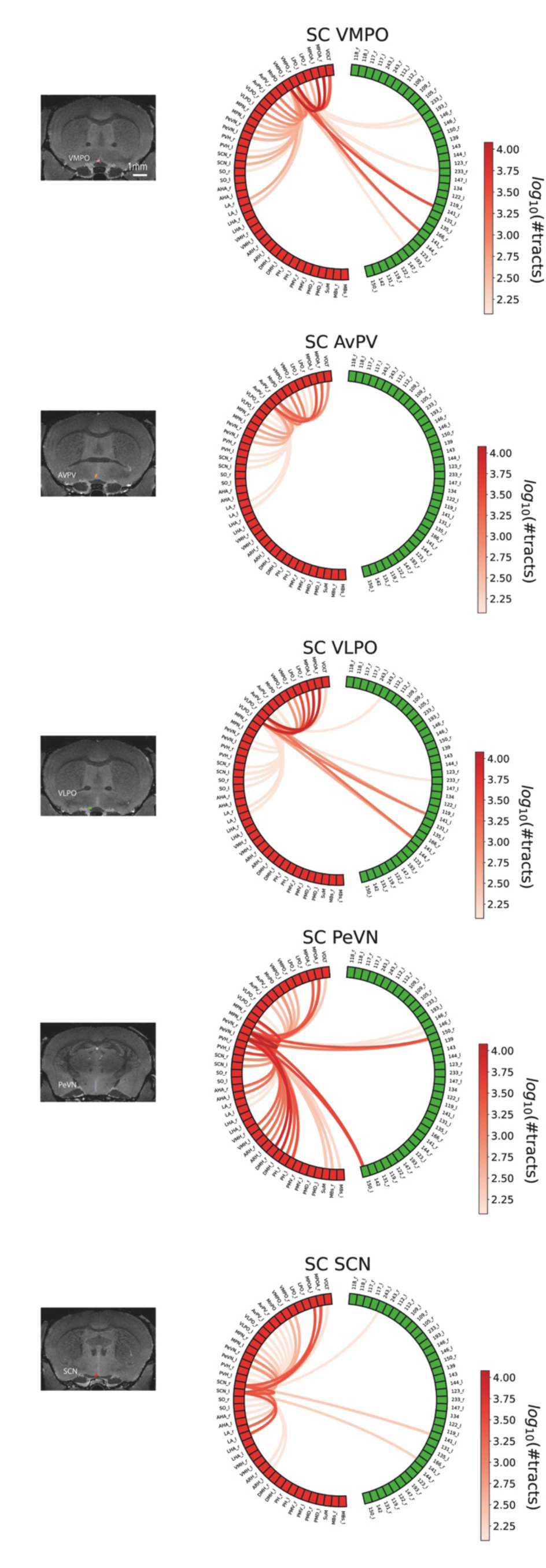
Structural Connectivity of Hypothalamic Nuclei Not Resolvable with rs-fMRI.

**Figure S2.**
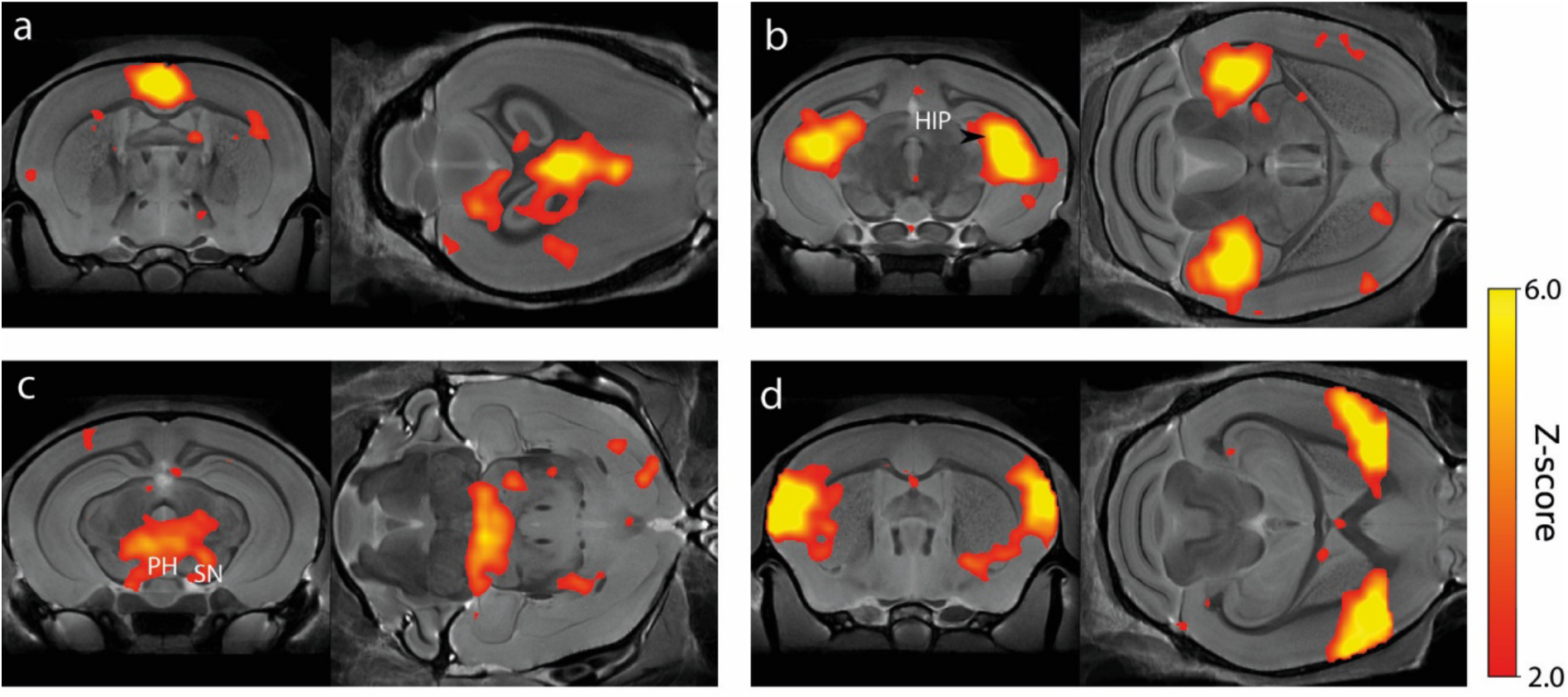
Independent component analysis (ICA) components in the mouse brain described previously by others^91, 92^. These components were found in the default mode network (a), the hippocampus (HIP) (b), the hypothalamus (c), and the lateral cortical network (d).

## References

1. Risold, P.Y., Thompson, R.H. & Swanson, L.W. The structural organization of connections between hypothalamus and cerebral cortex. Brain Res Brain Res Rev 24, 197–254 (1997).

2. Hahn, J.D., Sporns, O., Watts, A.G. & Swanson, L.W. Macroscale intrinsic network architecture of the hypothalamus. Proc Natl Acad Sci U S A 116, 8018–8027 (2019).

3. Fong, H., Zheng, J. & Kurrasch, D. The structural and functional complexity of the integrative hypothalamus. Science 382, 388–394 (2023).

4. Ahmed, R.M., Steyn, F. & Dupuis, L. Hypothalamus and weight loss in amyotrophic lateral sclerosis. Handb Clin Neurol 180, 327–338 (2021).

5. Stuber, G.D., Schwitzgebel, V.M. & Luscher, C. The neurobiology of overeating. Neuron 113, 1680–1693 (2025).

6. Kretzer, S. et al. The Dynamic Interplay Between Puberty and Structural Brain Development as a Predictor of Mental Health Difficulties in Adolescence: A Systematic Review. Biol Psychiatry 96, 585–603 (2024).

7. Mouse Genome Sequencing, C., et al. Initial sequencing and comparative analysis of the mouse genome. Nature 420, 520–562 (2002).

8. Lein, E.S. et al. Genome-wide atlas of gene expression in the adult mouse brain. Nature 445, 168–176 (2007).

9. Skarnes, W.C. et al. A conditional knockout resource for the genome-wide study of mouse gene function. Nature 474, 337–342 (2011).

10. Steuernagel, L. et al. HypoMap-a unified single-cell gene expression atlas of the murine hypothalamus. Nat Metab 4, 1402–1419 (2022).

11. Langlieb, J. et al. The molecular cytoarchitecture of the adult mouse brain. Nature 624, 333–342 (2023).

12. Moffitt, J.R. et al. Molecular, spatial, and functional single-cell profiling of the hypothalamic preoptic region. Science 362 (2018).

13. Bota, M., Dong, H.W. & Swanson, L.W. From gene networks to brain networks. Nat Neurosci 6, 795–799 (2003).

14. Simerly, R.B. in The Rat Nervous System (Fourth Edition). (ed. G. Paxinos) 267–294 (Academic press, 2015).

15. Thompson, R.H. & Swanson, L.W. Structural characterization of a hypothalamic visceromotor pattern generator network. Brain Res Brain Res Rev 41, 153–202 (2003).

16. Lanciego, J.L. & Wouterlood, F.G. Neuroanatomical tract-tracing techniques that did go viral. Brain Struct Funct 225, 1193–1224 (2020).

17. Xu, X. et al. Viral Vectors for Neural Circuit Mapping and Recent Advances in Trans-synaptic Anterograde Tracers. Neuron 107, 1029–1047 (2020).

18. Jiao, Z. et al. Projectome-based characterization of hypothalamic peptidergic neurons in male mice. Nat Neurosci 28, 1073–1088 (2025).

19. Johnson, G.A. et al. Waxholm space: an image-based reference for coordinating mouse brain research. Neuroimage 53, 365–372 (2010).

20. Baroncini, M. et al. MRI atlas of the human hypothalamus. Neuroimage 59, 168–180 (2012).

21. Florent, V. et al. Hypothalamic Structural and Functional Imbalances in Anorexia Nervosa. Neuroendocrinology 110, 552–562 (2020).

22. Ciobanu, L. et al. fMRI contrast at high and ultrahigh magnetic fields: insight from complementary methods. Neuroimage 113, 37–43 (2015).

23. Sazhina, T. et al. Time- and sex-dependent effects of juvenile social isolation on mouse brain morphology. Neuroimage 310, 121117 (2025).

24. Lopez-Kolkovsky, A.L., Meriaux, S. & Boumezbeur, F. Metabolite and macromolecule T1 and T2 relaxation times in the rat brain in vivo at 17.2T. Magn Reson Med 75, 503–514 (2016).

25. Dorr, A.E., Lerch, J.P., Spring, S., Kabani, N. & Henkelman, R.M. High resolution three-dimensional brain atlas using an average magnetic resonance image of 40 adult C57Bl/6J mice. Neuroimage 42, 60–69 (2008).

26. Ullmann, J.F., Watson, C., Janke, A.L., Kurniawan, N.D. & Reutens, D.C. A segmentation protocol and MRI atlas of the C57BL/6J mouse neocortex. Neuroimage 78, 196–203 (2013).

27. Richards, K. et al. Segmentation of the mouse hippocampal formation in magnetic resonance images. Neuroimage 58, 732–740 (2011).

28. Paxinos, G. & Franklin, K.B.J. The mouse brain in stereotaxic coordinates. (Academic Press, London; 2004).

29. Chung, S. et al. Identification of preoptic sleep neurons using retrograde labelling and gene profiling. Nature 545, 477–481 (2017).

30. Prager-Khoutorsky, M. & Bourque, C.W. Anatomical organization of the rat organum vasculosum laminae terminalis. Am J Physiol Regul Integr Comp Physiol 309, R324–337 (2015).

31. Gizowski, C. & Bourque, C.W. Neurons that drive and quench thirst. Science 357, 1092–1093 (2017).

32. Zimmerman, C.A., Leib, D.E. & Knight, Z.A. Neural circuits underlying thirst and fluid homeostasis. Nat Rev Neurosci 18, 459–469 (2017).

33. Machado, N.L.S. & Saper, C.B. Genetic identification of preoptic neurons that regulate body temperature in mice. Temperature (Austin) 9, 14–22 (2022).

34. Herbison, A.E. in Knobil and Neill’s Physiology of Reproduction, Edn. Fourth Edition. (eds. T.M. Plant & J. Zeleznik) pp 399-468 (Elsevier, New York; 2015).

35. Chachlaki, K., Garthwaite, J. & Prevot, V. The gentle art of saying NO: how nitric oxide gets things done in the hypothalamus. Nat Rev Endocrinol 13, 521–535 (2017).

36. Chachlaki, K. et al. NOS1 mutations cause hypogonadotropic hypogonadism with sensory and cognitive deficits that can be reversed in infantile mice. Sci Transl Med 14, eabh2369 (2022).

37. Chachlaki, K. et al. Phenotyping of nNOS neurons in the postnatal and adult female mouse hypothalamus. J Comp Neurol 525, 3177–3189 (2017).

38. Wu, Z., Autry, A.E., Bergan, J.F., Watabe-Uchida, M. & Dulac, C.G. Galanin neurons in the medial preoptic area govern parental behaviour. Nature 509, 325–330 (2014).

39. Barker, D.J. et al. Lateral preoptic area glutamate neurons relay nociceptive information to the ventral tegmental area. Cell Rep 42, 113029 (2023).

40. Sherin, J.E., Shiromani, P.J., McCarley, R.W. & Saper, C.B. Activation of ventrolateral preoptic neurons during sleep. Science 271, 216–219 (1996).

41. Liang, Y. et al. The NAergic locus coeruleus-ventrolateral preoptic area neural circuit mediates rapid arousal from sleep. Curr Biol 31, 3729–3742 e3725 (2021).

42. Tan, C.L. et al. Warm-Sensitive Neurons that Control Body Temperature. Cell 167, 47–59 e15 (2016).

43. Liu, D. et al. A hypothalamic circuit underlying the dynamic control of social homeostasis. Nature 640, 1000–1010 (2025).

44. Simerly, R.B. Wired on hormones: endocrine regulation of hypothalamic development. Curr.Opin.Neurobiol. 15, 81–85 (2005).

45. Waters, E.M. & Simerly, R.B. Estrogen induces caspase-dependent cell death during hypothalamic development. J Neurosci 29, 9714–9718 (2009).

46. Mittag, J. et al. Thyroid hormone is required for hypothalamic neurons regulating cardiovascular functions. J Clin Invest 123, 509–516 (2013).

47. Korchynska, S. et al. A hypothalamic dopamine locus for psychostimulant-induced hyperlocomotion in mice. Nat Commun 13, 5944 (2022).

48. Hastings, M.H., Maywood, E.S. & Brancaccio, M. Generation of circadian rhythms in the suprachiasmatic nucleus. Nat Rev Neurosci 19, 453–469 (2018).

49. Biag, J. et al. Cyto- and chemoarchitecture of the hypothalamic paraventricular nucleus in the C57BL/6J male mouse: a study of immunostaining and multiple fluorescent tract tracing. J Comp Neurol 520, 6–33 (2012).

50. Rossi, M.A. Control of energy homeostasis by the lateral hypothalamic area. Trends Neurosci 46, 738–749 (2023).

51. Al-Massadi, O. et al. Multifaceted actions of melanin-concentrating hormone on mammalian energy homeostasis. Nat Rev Endocrinol 17, 745–755 (2021).

52. Barbier, M., Croizier, S., Alvarez-Bolado, G. & Risold, P.Y. The distribution of Dlx1-2 and glutamic acid decarboxylase in the embryonic and adult hypothalamus reveals three differentiated LHA subdivisions in rodents. J Chem Neuroanat 121, 102089 (2022).

53. Prager-Khoutorsky, M. & Bourque, C.W. Mechanical basis of osmosensory transduction in magnocellular neurosecretory neurones of the rat supraoptic nucleus. J Neuroendocrinol 27, 507–515 (2015).

54. Leng, G. & Russell, J.A. The osmoresponsiveness of oxytocin and vasopressin neurones: Mechanisms, allostasis and evolution. J Neuroendocrinol 31, e12662 (2019).

55. Perkinson, M.R., Kim, J.S., Iremonger, K.J. & Brown, C.H. Visualising oxytocin neurone activity in vivo: The key to unlocking central regulation of parturition and lactation. J Neuroendocrinol 33, e13012 (2021).

56. Khodai, T. & Luckman, S.M. Ventromedial Nucleus of the Hypothalamus Neurons Under the Magnifying Glass. Endocrinology 162 (2021).

57. Douglass, A.M. et al. Acute and circadian feedforward regulation of agouti-related peptide hunger neurons. Cell Metab 37, 708–722 e705 (2025).

58. Francois, M. et al. Leptin receptor neurons in the dorsomedial hypothalamus require distinct neuronal subsets for thermogenesis and weight loss. Metabolism 163, 156100 (2025).

59. Aitken, T.J. et al. Negative feedback control of hypothalamic feeding circuits by the taste of food. Neuron 112, 3354–3370 e3355 (2024).

60. Kim, K.S. et al. GLP-1 increases preingestive satiation via hypothalamic circuits in mice and humans. Science 385, 438–446 (2024).

61. Simonds, S.E. et al. Leptin mediates the increase in blood pressure associated with obesity. Cell 159, 1404–1416 (2014).

62. Bruning, J.C. & Fenselau, H. Integrative neurocircuits that control metabolism and food intake. Science 381, eabl7398 (2023).

63. Nampoothiri, S., Nogueiras, R., Schwaninger, M. & Prevot, V. Glial cells as integrators of peripheral and central signals in the regulation of energy homeostasis. Nat Metab 4, 813–825 (2022).

64. Rodriguez-Cortes, B. et al. Tanycytes: bloodhounds of the metabolic brain. Trends Endocrinol Metab (2025).

65. Korf, H.W. & Moller, M. Arcuate nucleus, median eminence, and hypophysial pars tuberalis. Handb Clin Neurol 180, 227–251 (2021).

66. Cavalcante, J.C., da Silva, F.G., Saenz de Miera, C. & Elias, C.F. The ventral premammillary nucleus at the interface of environmental cues and social behaviors. Front Neurosci 19, 1589156 (2025).

67. Tseng, Y.T., Schaefke, B., Wei, P. & Wang, L. Defensive responses: behaviour, the brain and the body. Nat Rev Neurosci 24, 655–671 (2023).

68. Kowalczyk, T., Staszelis, A., Kazmierska-Grebowska, P., Tokarski, K. & Caban, B. The Role of the Posterior Hypothalamus in the Modulation and Production of Rhythmic Theta Oscillations. Neuroscience 470, 100–115 (2021).

69. Kesner, A.J. et al. Hypothalamic Supramammillary Control of Cognition and Motivation. J Neurosci 43, 7538–7546 (2023).

70. McNaughton, N. & Vann, S.D. Construction of complex memories via parallel distributed cortical-subcortical iterative integration. Trends Neurosci 45, 550–562 (2022).

71. Swaab, D.F. & Fliers, E. A sexually dimorphic nucleus in the human brain. Science 228, 1112–1115 (1985).

72. Byne, W. The medial preoptic and anterior hypothalamic regions of the rhesus monkey: cytoarchitectonic comparison with the human and evidence for sexual dimorphism. Brain Res 793, 346–350 (1998).

73. Swaab, D.F. Chapter 5 Sexually dimorphic nucleus of the preoptic area (SDN-POA) = intermediate nucleus = interstitial nucleus of the anterior hypothalamus (INAH-1) = preoptic nucleus. Handb Clin Neurol 79, 127–133 (2003).

74. Roselli, C.E., Larkin, K., Resko, J.A., Stellflug, J.N. & Stormshak, F. The volume of a sexually dimorphic nucleus in the ovine medial preoptic area/anterior hypothalamus varies with sexual partner preference. Endocrinology 145, 478–483 (2004).

75. Gorski, R.A., Gordon, J.H., Shryne, J.E. & Southam, A.M. Evidence for a morphological sex difference within the medial preoptic area of the rat brain. Brain Res 148, 333–346 (1978).

76. Panzica, G.C., Viglietti-Panzica, C. & Balthazart, J. The sexually dimorphic medial preoptic nucleus of quail: a key brain area mediating steroid action on male sexual behavior. Front Neuroendocrinol 17, 51–125 (1996).

77. Simerly, R.B., Swanson, L.W. & Gorski, R.A. The distribution of monoaminergic cells and fibers in a periventricular preoptic nucleus involved in the control of gonadotropin release: immunohistochemical evidence for a dopaminergic sexual dimorphism. Brain Res 330, 55–64 (1985).

78. Larriva-Sahd, J. Ultrastructural evidence of a sexual dimorphism in the neuropil of the medial preoptic nucleus of the rat: a quantitative study. Neuroendocrinology 54, 416–419 (1991).

79. Perrin, S., Garreau, S. & Rochette, M. in the European Conference on Spacecraft Structures, Materials and Mechanical Testing (ed. K. Fletcher) ESA SP-581 (Noordwijk, The Netherlands; 2005).

80. Benjamini, Y. & Yekutieli, D. The control of false discovery rate in multiple testing under dependency. The Annals of Statistics 29, 1165–1188 (2001).

81. Bonnavion, P., Mickelsen, L.E., Fujita, A., de Lecea, L. & Jackson, A.C. Hubs and spokes of the lateral hypothalamus: cell types, circuits and behaviour. J Physiol 594, 6443–6462 (2016).

82. Dong, H.W. & Swanson, L.W. Organization of axonal projections from the anterolateral area of the bed nuclei of the stria terminalis. J Comp Neurol 468, 277–298 (2004).

83. Barbier, M. et al. Projections from the dorsomedial division of the bed nucleus of the stria terminalis to hypothalamic nuclei in the mouse. J Comp Neurol 529, 929–956 (2021).

84. Goto, M., Canteras, N.S., Burns, G. & Swanson, L.W. Projections from the subfornical region of the lateral hypothalamic area. J Comp Neurol 493, 412–438 (2005).

85. Canteras, N.S., Simerly, R.B. & Swanson, L.W. Organization of projections from the ventromedial nucleus of the hypothalamus: a Phaseolus vulgaris-leucoagglutinin study in the rat. J Comp Neurol 348, 41–79 (1994).

86. Bouret, S.G., Draper, S.J. & Simerly, R.B. Formation of projection pathways from the arcuate nucleus of the hypothalamus to hypothalamic regions implicated in the neural control of feeding behavior in mice. J.Neurosci. 24, 2797–2805 (2004).

87. van den Pol, A.N. & Cassidy, J.R. The hypothalamic arcuate nucleus of rat--a quantitative Golgi analysis. J Comp Neurol 204, 65–98 (1982).

88. Prevot, V. et al. The Versatile Tanycyte: A Hypothalamic Integrator of Reproduction and Energy Metabolism. Endocr Rev 39, 333–368 (2018).

89. Ugurbil, K. Imaging at ultrahigh magnetic fields: History, challenges, and solutions. Neuroimage 168, 7–32 (2018).

90. Beckmann, C.F. & Smith, S.M. Probabilistic independent component analysis for functional magnetic resonance imaging. IEEE Trans Med Imaging 23, 137–152 (2004).

91. Zerbi, V., Grandjean, J., Rudin, M. & Wenderoth, N. Mapping the mouse brain with rs-fMRI: An optimized pipeline for functional network identification. Neuroimage 123, 11–21 (2015).

92. Liska, A., Galbusera, A., Schwarz, A.J. & Gozzi, A. Functional connectivity hubs of the mouse brain. Neuroimage 115, 281–291 (2015).

93. Andersson, J.L., Skare, S. & Ashburner, J. How to correct susceptibility distortions in spin-echo echo-planar images: application to diffusion tensor imaging. Neuroimage 20, 870–888 (2003).

94. Smith, S.M. et al. Advances in functional and structural MR image analysis and implementation as FSL. Neuroimage 23 Suppl 1, S208–219 (2004).

95. Beckmann, C.F., Mackay, C.E., Filippini, N. & Smith, S.M. Group comparison of resting-state FMRI data using multi-subject ICA and dual regression. Neuroimage 47, S148 (2009).

96. Canteras, N.S. & Swanson, L.W. Projections of the ventral subiculum to the amygdala, septum, and hypothalamus: a PHAL anterograde tract-tracing study in the rat. J Comp Neurol 324, 180–194 (1992).

97. Thompson, R.H. & Swanson, L.W. Organization of inputs to the dorsomedial nucleus of the hypothalamus: a reexamination with Fluorogold and PHAL in the rat. Brain Res Brain Res Rev 27, 89–118 (1998).

98. Sternson, S.M., Shepherd, G.M. & Friedman, J.M. Topographic mapping of VMH --> arcuate nucleus microcircuits and their reorganization by fasting. Nat Neurosci 8, 1356–1363 (2005).

99. Fernandois, D. et al. Estrogen receptor-alpha signaling in tanycytes lies at the crossroads of fertility and metabolism. Metabolism, 155976 (2024).

100. Lhomme, T. et al. Tanycytic networks mediate energy balance by feeding lactate to glucose-insensitive POMC neurons. J Clin Invest 131, e140521 (2021).

101. Mullier, A., Bouret, S.G., Prevot, V. & Dehouck, B. Differential distribution of tight junction proteins suggests a role for tanycytes in blood-hypothalamus barrier regulation in the adult mouse brain. J Comp Neurol 518, 943–962 (2010).

102. Langlet, F. et al. Tanycytic VEGF-A Boosts Blood-Hypothalamus Barrier Plasticity and Access of Metabolic Signals to the Arcuate Nucleus in Response to Fasting. Cell Metab 17, 607–617 (2013).

103. Lei, H., Poitry-Yamate, C., Preitner, F., Thorens, B. & Gruetter, R. Neurochemical profile of the mouse hypothalamus using in vivo 1H MRS at 14.1T. NMR Biomed 23, 578–583 (2010).

104. Urenjak, J., Williams, S.R., Gadian, D.G. & Noble, M. Proton nuclear magnetic resonance spectroscopy unambiguously identifies different neural cell types. J Neurosci 13, 981–989 (1993).

105. Isaacks, R.E., Bender, A.S., Kim, C.Y., Prieto, N.M. & Norenberg, M.D. Osmotic regulation of myo-inositol uptake in primary astrocyte cultures. Neurochem Res 19, 331–338 (1994).

106. Rothermundt, M. et al. Glial cell activation in a subgroup of patients with schizophrenia indicated by increased S100B serum concentrations and elevated myo-inositol. Prog Neuropsychopharmacol Biol Psychiatry 31, 361–364 (2007).

107. Kim, H., Choi, S., Lee, E., Koh, W. & Lee, C.J. Tonic NMDA Receptor Currents in the Brain: Regulation and Cognitive Functions. Biol Psychiatry 96, 164–175 (2024).

108. Amunts, K. et al. The Human Brain Project: Creating a European Research Infrastructure to Decode the Human Brain. Neuron 92, 574–581 (2016).

109. Amunts, K. et al. The Human Brain Project-Synergy between neuroscience, computing, informatics, and brain-inspired technologies. PLoS Biol 17, e3000344 (2019).

110. Tkac, I. et al. Regional sex differences in neurochemical profiles of healthy mice measured by magnetic resonance spectroscopy at 9.4 tesla. Front Neurosci 17, 1278828 (2023).

111. Gaudin, R., Bernard, J., Glatigny, M. & Boido, D. A 3D-printed cradle for mouse preclinical MRI with an integrated water heating system. bioRxiv, 10.1101/2024.1112.1129.630663 (2024).

112. Garwood, M. & DelaBarre, L. The return of the frequency sweep: designing adiabatic pulses for contemporary NMR. J Magn Reson 153, 155–177 (2001).

113. Tkac, I., Starcuk, Z., Choi, I.Y. & Gruetter, R. In vivo 1H NMR spectroscopy of rat brain at 1 ms echo time. Magn Reson Med 41, 649–656 (1999).

114. Avants, B.B. et al. A reproducible evaluation of ANTs similarity metric performance in brain image registration. Neuroimage 54, 2033–2044 (2011).

115. Jenkinson, M., Beckmann, C.F., Behrens, T.E., Woolrich, M.W. & Smith, S.M. Fsl. Neuroimage 62, 782–790 (2012).

116. Yushkevich, P.A. et al. User-guided 3D active contour segmentation of anatomical structures: significantly improved efficiency and reliability. Neuroimage 31, 1116–1128 (2006).

117. Lin, Y. et al. RS(2)-Net: An end-to-end deep learning framework for rodent skull stripping in multi-center brain MRI. Neuroimage 298, 120769 (2024).

118. Buades, A., Coll, B. & Morel, J.-M. Non-Local Means Denoising. Image Processing On Line 208–212 (2011).

119. Basser, P.J., Mattiello, J. & LeBihan, D. MR diffusion tensor spectroscopy and imaging. Biophys J 66, 259–267 (1994).

120. Descoteaux, M., Angelino, E., Fitzgibbons, S. & Deriche, R. Regularized, fast, and robust analytical Q-ball imaging. Magn Reson Med 58, 497–510 (2007).

121. Lee, S.-H., Ban, W. & Shih, Y.-Y.I. BrkRaw/bruker : BrkRaw v0.3.3 version 0.3.3. zenodo, 10.5281/zenodo.3877179 (2020).

122. Cox, R.W. AFNI : software for analysis and visualization of functional magnetic resonance neuroimages. Computers and Biomedical research 29, 162–173 (1996).

123. Cox, R.W. & Hyde, J.S. Software tools for analysis and visualization of fMRI data. NMR Biomed 10, 171–178 (1997).

124. Tustison, N.J. et al. The ANTsX ecosystem for quantitative biological and medical imaging. Sci Rep 11, 9068 (2021).

125. Desrosiers-Gregoire, G., Devenyi, G.A., Grandjean, J. & Chakravarty, M.M. A standardized image processing and data quality platform for rodent fMRI. Nat Commun 15, 6708 (2024).

126. Wang, L. et al. fMRI signals in white matter rewire gray matter community organization. Neuroimage 297, 120763 (2024).

127. Schilling, K.G. et al. Whole-brain, gray, and white matter time-locked functional signal changes with simple tasks and model-free analysis. Proc Natl Acad Sci U S A 120, e2219666120 (2023).

128. Zalesky, A., Fornito, A. & Bullmore, E.T. Network-based statistic: identifying differences in brain networks. Neuroimage 53, 1197–1207 (2010).

129. Pijnappel, W.W.F., van den Boogaart, A., de Beer, R. & van Ormondt, D. SVD-based quantification of magnetic resonance signals. Journal of Magnetic Resonance (1969) 97, 122–134 (1992).

130. Provencher, S.W. Estimation of metabolite concentrations from localized in vivo proton NMR spectra. Magn Reson Med 30, 672–679 (1993).

